# Expression of clock genes tracks daily and tidal time in brains of intertidal crustaceans *Eurydice pulchra* and *Parhyale hawaiensis*

**DOI:** 10.1101/2025.01.13.632323

**Authors:** A. Oliphant, C.Y. Sia, C.P. Kyriacou, D.C. Wilcockson, M.H. Hastings

## Abstract

Intertidal organisms, such as the crustaceans *Eurydice pulchra* and *Parhyale hawaiensis*, express daily and tidal rhythms of physiology and behaviour to adapt to their temporally complex environments. Although the molecular-genetic basis of the circadian clocks driving daily rhythms in terrestrial animals is well understood, the nature and mechanism of the circatidal clocks driving tidal rhythms remain a mystery. Using *in situ* hybridisation, we identified discrete clusters of ∼60 putative “clock” cells co-expressing canonical circadian clock genes with comparable distributions across the protocerebrum of *E. pulchra* and *P. hawaiensis* brains. In tidally rhythmic, field-collected *E. pulchra* sampled under a light:dark (LD) cycle, the expression of *period* (*per*) and *cryptochrome 2* (*cry2*) exhibited daily rhythms in particular cell groups (e.g., medioposterior cells) whereas *timeless* (*tim*) showed 12-hour rhythms in others (e.g., medial cells). In tidally rhythmic laboratory-reared *P. hawaiensis*, previously entrained to 12.4-hour cycles of agitation under LD and sampled under continuous darkness, several cell groups (e.g., medioposterior cells) exhibited circadian expression of *per* and *cry2*. In contrast, dorsal-lateral cells in the protocerebrum exhibited robust ∼12-hour, i.e., circatidal, rhythms of *per* and *cry2*, phased to the prior tidal agitation but not the prior LD. In *P. hawaiensis* exhibiting daily behaviour under LD without tidal agitation, robust daily rhythms of *per* and *cry2* expression were evident in medioposterior and other cells whereas expression in dorsal-lateral cells was not rhythmic, underlining their intrinsic tidal periodicity. These results implicate canonical circadian mechanisms in circatidal time-keeping and reveal conserved brain networks as potential neural substrates for the generation of interactive daily and tidal rhythms appropriate to intertidal habitats.

## Introduction

Organisms adapt to their cyclical environments through rhythms of metabolism, physiology and behaviour that are driven by internal biological clocks ^1^. Terrestrial environmental cycles are predominantly daily, leading to the evolution of clocks with an autonomous period of approximately one day (hence *circa*-*dian*) and entrained to the cycle of light and darkness. In contrast, coastal environments present greater temporal complexity and marine organisms have additionally evolved a range of circatidal (∼12.4 hours), circasemilunar (∼14.8 days) and circalunar (∼29.5 days) timing mechanisms ^2 3^ that co-ordinate adaptive rhythms across these time domains. Despite their biological relevance, the mechanistic underpinnings of circatidal clocks remain a mystery and three principal hypotheses have been proposed ^2^. First, circatidal and circadian rhythms are driven by a single oscillatory system the period of which is plastic, adapting to the period of the dominant environmental cue ^4^. Second, paired circalunidian (∼24.8-hour, i.e., slightly modified circadian) oscillators run in antiphase to create 12.4-hour rhythms ^5^. Third, separate 12.4-hour and 24-hour oscillators co-exist, independently driving circatidal and daily phenotypes ^6^. Testing these models requires identification of the clock cells and genes responsible for tidal time-keeping.

Circadian clocks in animals pivot around a transcriptional/ translational feedback loop (TTFL) in which positive factors CLOCK (CLK) and BMAL1 (or CYC in flies), acting through E-box promotor sites, transactivate genes that encode negative regulators, PERIOD (PER), CRYPTOCHROME2 (CRY2) and/ or TIMELESS (TIM), depending on species ^7^. Accumulating negative regulators progressively suppress transactivation and following their degradation the cycle restarts after ∼24 hours ^8 9^. The intertidal isopod *Eurydice pulchra* and amphipod *Parhyale hawaiensis* can express autonomous circadian and/ or circatidal rhythms of locomotor behaviour in the absence of environmental cues ^10 11 12 13^. However, the only molecular markers of circatidal timekeeping relate to mitochondrial gene expression and cellular redox state in *E. pulchra* ^14^. Nevertheless, genes encoding CLOCK, BMAL1, PER and CRY2 are present in both species ^12 15^ and *tim* expression in the head of *E. pulchra* is circadian ^12 16^ (*P. hawaiensis* lacks *tim* ^15^). *E. pulchra* CLOCK:BMAL1 heterodimers can transactivate E-boxes and are suppressed by PER and CRY2 ^12^, whilst PhBMAL1 co-expressed with mouse CLK can also transactivate E-boxes ^13^. Finally, genomic deletion (*P. hawaiensis*) or RNAi-knock-down (*E. pulchra*) of *Bmal1* compromises tidal activity rhythms^13 16^, raising the intriguing possibility of interactions between circadian and circatidal clocks at the molecular-genetic level.

The current aim was to describe the expression of *Bmal1*, *Clk*, *per* and *cry2* (together with *Eptim*) across the brain to identify putative “clock” cells, and then to test their relationship to daily, circadian and circatidal behaviour. We reveal ∼60 such cells arranged in multiple, discrete cell groups in the brain and show that particular groups are capable of tracking daily/ circadian time, encoded as the rhythmic expression of canonical circadian genes. Furthermore, we also identify cells with 12-hour rhythms of expression of *tim* in *E. pulchra* and of *per* and *cry2* in *P. hawaiensis* and show that the latter are capable of tracking circatidal time. These results implicate canonical circadian mechanisms in circatidal time-keeping. They also reveal brain networks that may be neural substrates for the generation of interactive daily and tidal rhythms appropriate to local intertidal habitats.

## Results

### Cellular expression of circadian clock genes in the brains of *E. pulchra* and *P. hawaiensis*

To map gene expression across the brain, we first generated reference brains for *E. pulchra* and *P. hawaiensis* by averaging confocal image stacks of brains immunolabelled with anti-SYNORF1 (Supplementary Figure 1). We segmented the neuropils to facilitate description of gross brain anatomy and naming of clock cell groups relative to the body axis (Supplementary Figure 2, Supplementary video 1, 2). Our neuroanatomical descriptions of *E. pulchra* are based on the isopod *Saduria entomon* ^17^. The brain structure of *P. hawaiensis* has been described ^18 19^.

*Bmal1* is implicated in circadian and circatidal timekeeping in both *E. pulchra* and *P. hawaiensis*. We therefore used whole-mount HCR-FISH to map *Bmal1* transcripts as a first step to identify putative “clock” cells. This revealed widespread expression across the brain of *E. pulchra* (Supplementary Figure 3A) and, consistent with qPCR studies ^12^, its intensity did not change over time in LD (Supplementary Figure 4A). The expression of *Clk* mRNA was similarly widespread in the brain of *E. pulchra*, with most cells expressing low levels although there was a condensation of higher intensity signals in ∼60 cells, located principally in the median protocerebrum (Supplementary Figure 3B). These *Bmal1* and *Clk* expression patterns in *E. pulchra* were confirmed using RNAscope^TM^ on brain sections (Supplementary Figure 5).

Pan-cellular expression of *Bmal1* was also evident in the brain of *P. hawaiensis* (Supplementary Figure 3C) and, again, this did not change across the light:dark (LD) cycle (Supplementary Figure 4B). In contrast, the expression of *Clk* in *P. hawaiensis* was less widespread than in *E. pulchra*, with enrichment in cells in the median protocerebrum (Supplementary Figure 3D). The enrichment was clearer than in *E. pulchra* and these enriched cells were comparable in number and location to those in *E. pulchra*. Thus, notwithstanding the requirement for *Bmal1* to sustain circatidal behaviour in both species ^13 16^, the pattern of its expression did not specifically signpost any populations of putative clock cells. In contrast, the expression of *Clk* was significantly elevated in discrete cell populations in the median protocerebrum of both species.

We then mapped the expression of genes encoding negative regulators: *per*, *cry2* and *tim* across the brain of *E. pulchra*. This revealed strong, specific signal for all three transcripts restricted to discrete cell clusters in the median protocerebrum (Supplementary Figure 3E, F, G). In *P. hawaiensis* brain the expression of *per* and *cry2* exhibited a pattern similar to that of *E. pulchra*, with high levels of both transcripts localised to ∼60 cells in the median protocerebrum (Supplementary Figure 3I, J). The HCR-FISH was specific, insofar as probes against *deGFP* gave no signal Supplementary Figure 3H, K).

### Co-expression of circadian clock genes in distinct cell groups of *E. pulchra* and *P. hawaiensis* brains

CLOCK:BMAL1 heterodimers are required for transactivation within the TTFL and so we compared the co-expression of *Bmal1* and *Clk* by multiplexed HCR-FISH of *E. pulchra* brains dissected in the light phase. The *Clk*-enriched cells expressed *Bmal1* at the low level seen in other cells across the brain. They also expressed high levels of *per* (Figure 1A) and the *per*-expressing cells also expressed *cry2* and *tim* (Figure 1B, C). Given that *per* and *cry2* encode the principal negative regulators of E-box-mediated transcription in *E. pulchra* ^12^ this small population of cells in the median protocerebrum co-expressed the full complement of genes required for a functional TTFL, highlighting them as putative clock cells. Using a similar approach in *P. hawaiensis* we found, again, that *Clk*-enriched cells expressed *Bmal1* and high levels of *cry2* (Figure 1D). Furthermore, the *cry2*-enriched (and hence, *Clk*-enriched) cells also expressed *per* (Figure 1E), indicating that they are likely TTFL-competent cells. By combining HCR-FISH with anti-SYNORF1 immunofluorescence (Supplementary Figure 6, Supplementary video 1, 2) the putative clock cell clusters were registered to the reference brains (Figure 2). The groups were named by general location: 8 in *E. pulchra* and 9 in *P. hawaiensis*, with a total of ∼30 cells per hemisphere, distributed symmetrically (Supplementary Table 1, Supplementary Results).

**Figure 1.**
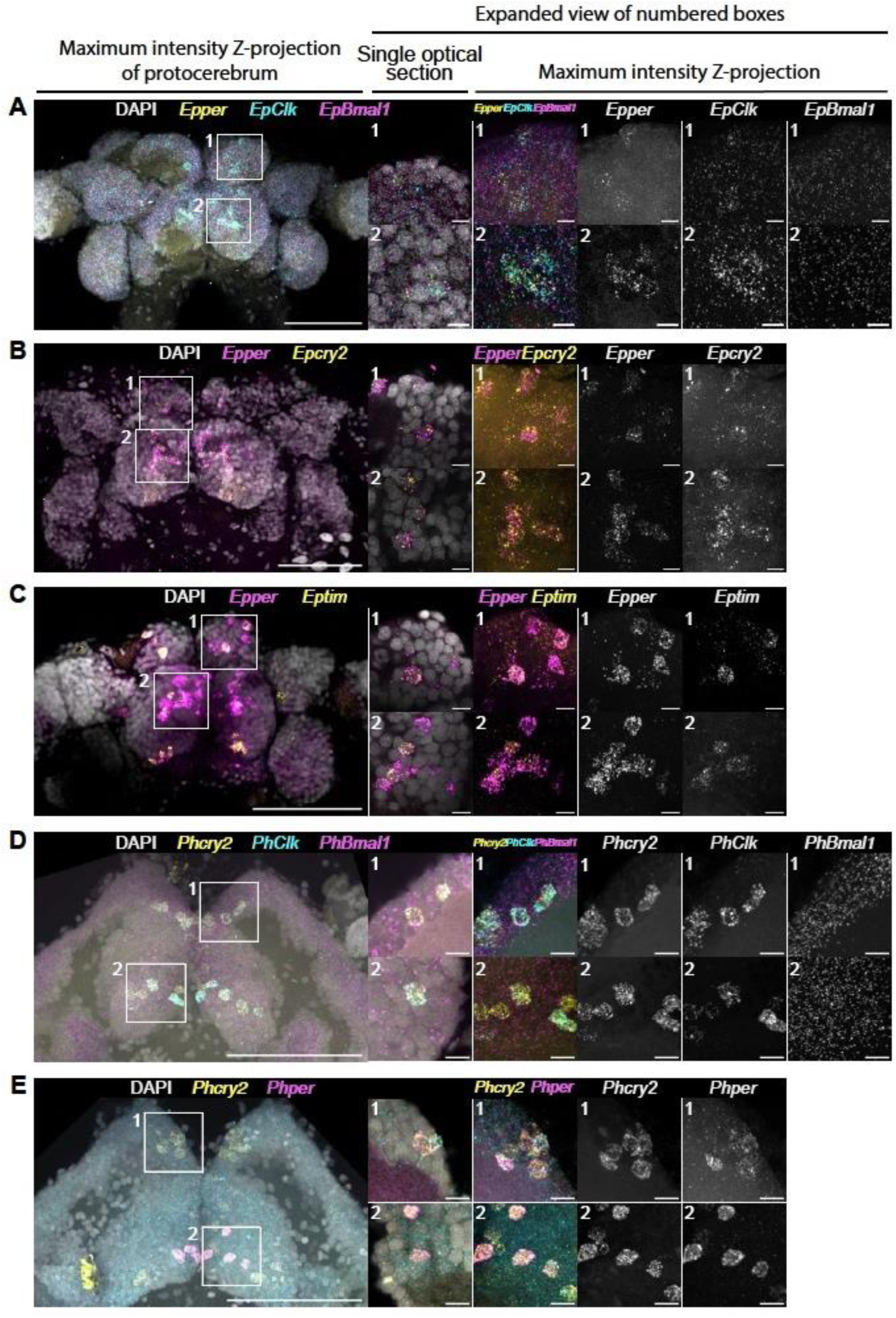
Cellular co-expression of circadian clock genes in the brains of *E. pulchra* and *P. hawaiensis*. (A) Representative maximum intensity Z-projections and single optical sections of *E. pulchra* brain probed for *Epper*, *EpClk* and *EpBmal1* using HCR-FISH. Left: merged low-power (25x) view of protocerebrum of *E. pulchra*, centre: merged high-power (40x) single optical sections in boxes labelled 1 and 2, right: single channel high-power maximum intensity Z-projections. *EpBmal1* is widely and uniformly expressed in both putative clock cells and other cells across the brain, but it is not enriched in cells enriched for *EpClk*, whereas *Epper* is enriched in *EpClk*-enriched cells. (B) As in A for *Epper* and *Epcry2*, expression of which is co-enriched in putative clock cells but not in other cells in the brain of *E. pulchra*. (C) As in A for *Epper* and *Eptim*, of *Epper*-enriched cells in *E. pulchra*. (D) As in A but for *P. hawaiensis* brain probed for *PhBmal1*, *PhClk* and *Phcry2* using HCR-FISH. *PhBmal1* is widely and uniformly expressed in both putative clock cells and other cells across the brain, but it is not enriched in cells enriched for *PhClk*, whereas *Phcry2* is enriched in *PhClk*-enriched cells. (E) As in A but for *P. hawaiensis* brain labelled for *Phcry2* and *Phper*. *Phcry2* and *Phper* are co-enriched in putative clock cells, but not other cells in the brain. Scale bars: 100 µm (left-most panels), 10 µm (other panels).

**Figure 2.**
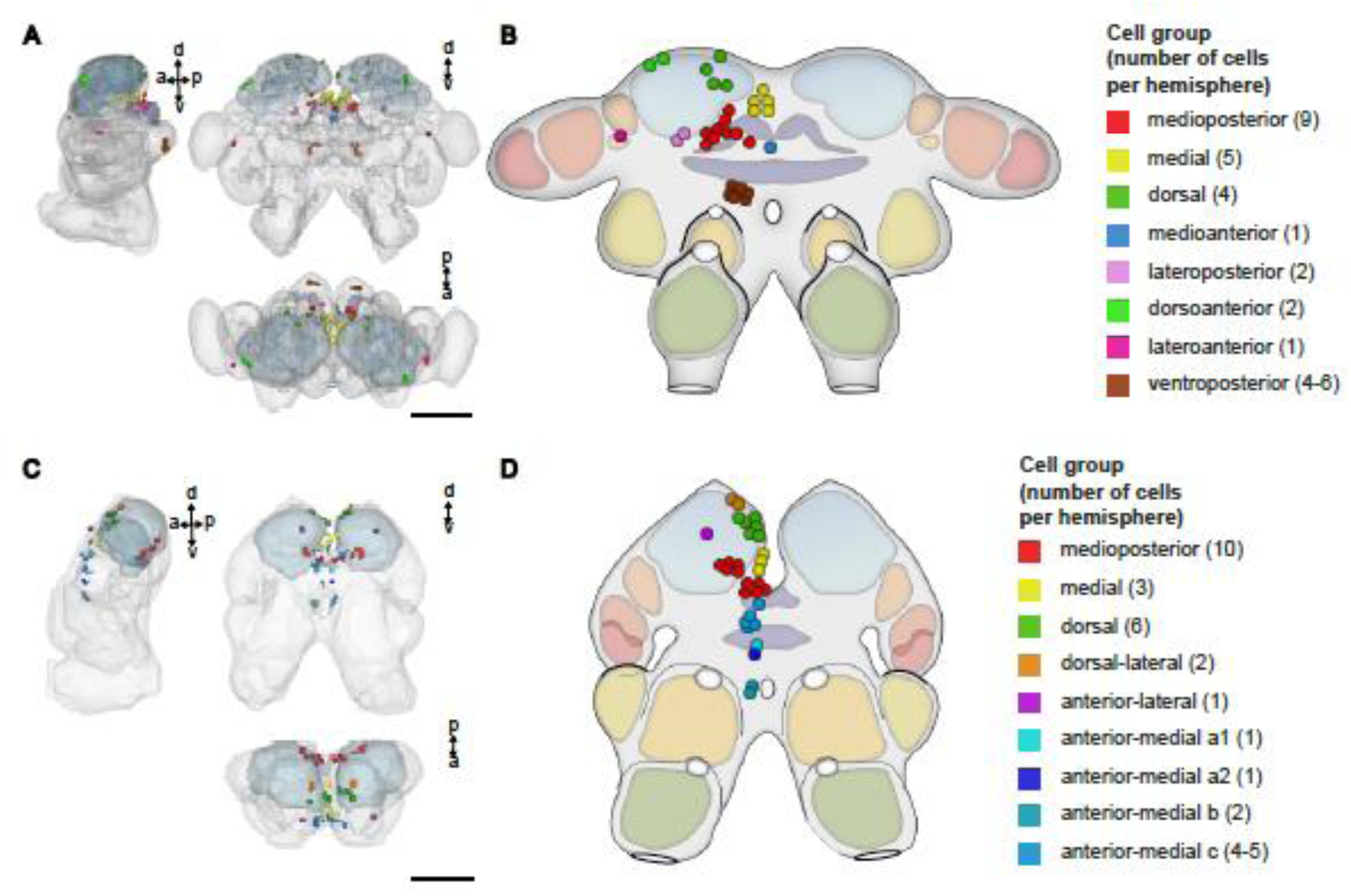
Putative clock cells in the brains of *E. pulchra* and *P. hawaiensis* revealed by co-expression of canonical circadian clock genes. (A) Lateral, frontal and dorsal views of the *E. pulchra* reference brain model (from Supplementary Figure 2A) on to which somata enriched for expression of *Epper* and *Epcry2* are registered and segmented. The neuropils referred to in the Supplementary Materials (Supplementary Results) to signpost putative clock cells are here made translucent. (B) Diagrammatic representation of the *E. pulchra* brain summarising the locations of cells co-enriched for *Epper* and *Epcry2*, based on the 3D reconstruction in A. (C) Lateral, frontal and dorsal views of the *P. hawaiensis* reference brain model (from Supplementary Figure 2C) on to which somata enriched for expression of *Phper* and *Phcry2* are registered and segmented. The neuropils referred to in the Supplementary Materials (Supplementary Results) to signpost putative clock cells are here made translucent. (D) Diagrammatic representation of the *P. hawaiensis* brain summarising the locations of cells co-enriched for *Phper* and *Phcry2*, based on the 3D reconstruction in C. Compass markers in A and C show anterior (a), posterior (p), dorsal (d) and ventral (v) directions. Discrete cell groups are colour coded (see key). Cell numbers within each group (per hemisphere) in parenthesis. Scale bars: 100 µm.

### Rhythmic expression of circadian clock genes in circatidally rhythmic *E. pulchra* under a light:dark cycle

When placed into free-running conditions (DD) in the laboratory, field-collected *E pulchra* displayed strong circatidal activity rhythms, with peak activity corresponding to the expected time of high water on the home beach (Supplementary Figure 7). Across 11 independent field collections, 54.5 ±6.5% of animals were tidally rhythmic (range: 20–76%) with a mean period of 12.36 ±0.02 h (±SEM, N =478). To test whether some or all putative clock cells of *E. pulchra* can track daily or tidal time, we performed whole-mount, multiplexed HCR-FISH on dissected brains of circatidally rhythmic beach-collected *E. pulchra* sampled every 2 hours under LD (16:8) for 24 hours (Figure 3A). The number of HCR-FISH spots was used to quantify transcript abundance, and we considered expression to be rhythmic only when Kruskal-Wallis/ ANOVA identified a significant time effect and both JTK_CYC ^20^ and RAIN ^21^ were significant (p<0.05) at a period of 12 or 24 hours, for circatidal and daily rhythms, respectively. The aggregated counts across all cells, as a measure of total brain expression, showed strong daily rhythms for *per*, *cry2* and *tim*, with peak levels in the late dark phase (Figure 3B). Aggregate levels of *tim* also showed a statistically significant 12-hour rhythm, although the overall profiles for aggregate *per*, *cry2* and *tim* were similar and tracked daily time.

**Figure 3.**
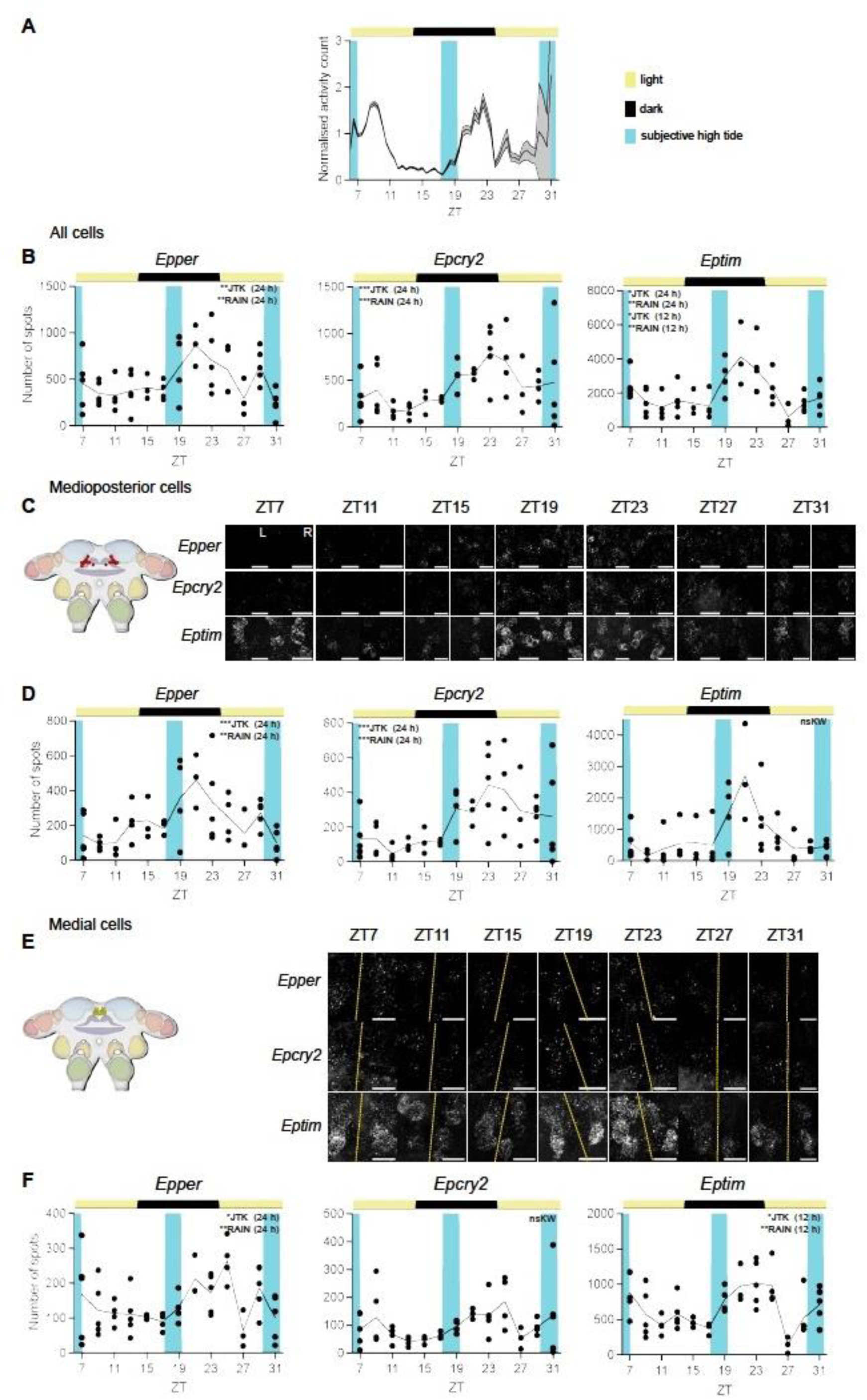
Time course of expression of *per, cry2 and tim* in cells of tidally rhythmic *E. pulchra* held on a LD cycle. (A) Normalised swimming activity (mean +SEM, shading) of beach-collected *E. pulchra* harvested under LD (16L:8D) for the HCR-FISH time-course (initial group size at start of harvest at ZT7 =65). (B). Mean aggregate expression, quantified as the number of FISH spots, of *Epper, Epcry2* and *Eptim* across all cells plotted across time in beach-collected animals held under LD. Each point is from one brain. (C) Left: cartoon to show location of medioposterior cell group in the brain. Right: representative maximum intensity Z-projections of *Epper*, *Epcry2* and *Eptim* expression in both hemispheres (L: left, R: right) across ZT (individual HCR-FISH channels in rows). Scale bars: 20 µm. (D) Mean expression, quantified as the number of FISH spots, of *Epper* and *Epcry2* in medioposterior cell group plotted across time in beach-collected animals held under LD. Each point is from one brain. (E, F) As C and D but for medial cells group. Yellow dashed lines: midlines. Scale bars: 10 µm. Inset text indicates statistical tests (JTK-cycle and RAIN) performed to determine rhythmicity following significant time effect by ANOVA or Kruskall-Wallis. *p* <0.001***, *p* <0.01**, *p* ≤0.05*, *p* >0.05^ns^. See also Supplementary Figure 8 for clock gene expression in lateroposterior cell group. For statistics and data shown in this figure, refer to Supplementary Table 2.

To explore whether circatidal and circadian behavioural phenotypes are generated by different cell populations that contain ∼12-hour and ∼24-hour molecular timers, we analysed clock gene expression at the level of individual cell groups. Of the 7 examined, 5 showed significant rhythms of one or multiple transcripts (Supplementary Table 2). Daily patterning was evident for *per* and *cry2* expression in the medioposterior cells with peaks in late night (Figure 3C, D), whilst medial cells had a daily rhythm of *per* but a 12-hour rhythm of *tim* (Figure 3E, F). Lateroposterior cells also had a significant 12-hour cycle of *tim* (Supplementary Figure 8B), whereas *per* and *cry2* levels in these cells were not significantly rhythmic. In summary, in circatidally behaving *E. pulchra* sampled under LD, aggregate expression of *per* and *cry2* tracked daily time, whilst aggregate levels of *tim* were more complex, reflecting both daily and circatidal time. When examined across distinct cell groups, daily cycles of *per* and *cry2* were evident in 4 of 7 groups whilst a 12-hour, putative circatidal, pattern of *tim* expression was detected in two. Thus, distinct cell groups in *E. pulchra* appear able to track either daily or tidal time.

### Rhythmic expression of circadian clock genes in circatidally active *P. hawaiensis*

To establish circatidal behavioural rhythms, *P. hawaiensis* laboratory-reared under a 12L:12D cycle were entrained by a cycle of mechanical agitation for 2 hours every 12.4 hours for >30 days. On transfer to free-running conditions of DD and no further agitation, they showed circatidal rhythms with peak activity phased to anticipated agitation (i.e., subjective high-tide) (Supplementary Figure 9A, B). Across 13 experiments, 52.8 ±2.8% of animals showed circatidal behaviour (range 40 –70%) with a mean period of 12.40 ±0.02 h (±SEM, N =456) (Supplementary Figure 9C, D). To confirm that the circatidal behaviour was phased by the agitation and not by the LD cycle, a sub-group of animals were subjected to a phase-shifted regime for the final 6 days of entrainment in which agitation occurred every 12.9 hours rather than every 12.4 hours as in matched controls. It therefore accumulated a relative delay of ∼6 hours (0.5 hours per cycle over ∼12 cycles). On release to DD, control and phase-shifted animals exhibited circatidal behavioural rhythms with peak activity at projected high tide and periods of ∼12.4 h that were not significantly different between groups (Kolmogorov-Smirnov test: *D* =0.1779, p =0.59, n.s) (Supplementary Figure 9E, F, G). Over four replications of the control and shifted regimes, their effectiveness (45.7% vs. 49.8% circatidal animals) was comparable (paired t-test: t =0.27, df =3, p =0.80, n.s., Cohen’s d =0.136). Importantly, the phase of the circatidal behaviour was delayed by 4.94 ±0.51 h (±SEM, N =4) in phase-shifted animals, relative to the controls (Supplementary Figure 9E, F, H). This not only confirms the capacity of *P. hawaiensis* to exhibit circatidal behaviour ^13^ but it also demonstrates that the circatidal clockwork of *P. hawaiensis* is sensitive to mechanical agitation, as is the case for *E. pulchra* ^11^.

To test whether cells in *P. hawaiensis* could track circadian and/ or tidal time, whole-mount HCR-FISH was applied to the brains of animals entrained by tidal agitation and exhibiting circatidal activity before harvest (Figure 4A). Aggregate expression of *per* was circadian, peaking in subjective night, whilst aggregate expression of *cry2* did not vary with time (Figure 4B). Across the 10 cell groups examined, 3 showed significant rhythms of one or multiple transcripts (Supplementary Table 3). Notably, expression of *per* was strongly circadian in medioposterior cells and anterior-medial a1 cells (Figure 4C, D and E, F). Moreover, these cells also showed circadian cycles of *cry2* expression, running in antiphase to *per*. Thus, *per* and *cry2* expression in these cells tracked circadian time rather than tidal time in circatidally rhythmic animals. In contrast, dorsal-lateral cells in the same animals showed a significant 12-hour cycle of *per* and *cry2* transcripts (Figure 4G, H). Furthermore, in the dorsal-lateral cells, unlike the circadian cells, *per* and *cry2* oscillated almost in phase, both peaking after subjective high tide (∼CT00-03 and CT15).

**Figure 4.**
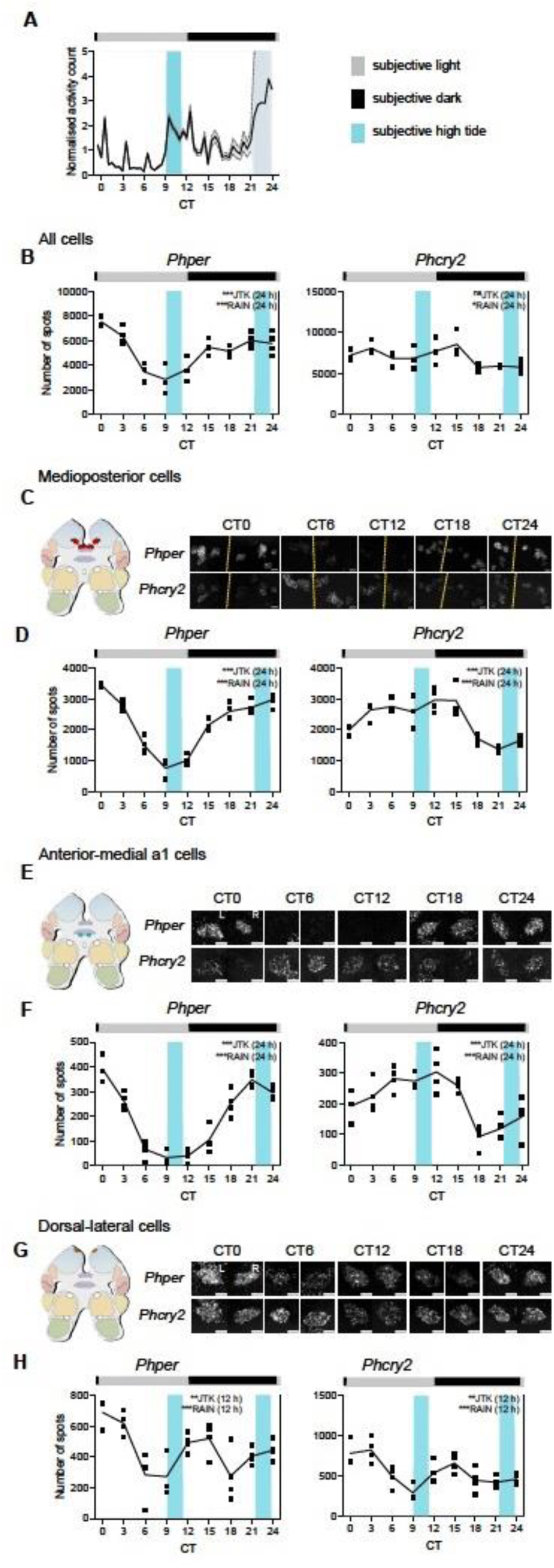
Free-running clock gene expression rhythms in tidally entrained *P. hawaiensis* (tidal phase θ). (A) Normalised swimming activity (mean +SEM, shading) of *P. hawaiensis* synchronised to the prior agitation cycle and harvested for the HCR-FISH time-course (initial group size at start of harvest at CT0 =37). (B) Mean aggregate expression, quantified as the number of FISH spots, of *Phper* and *Phcry2* across all putative clock cells plotted across time in tidally entrained animals sampled under DD. Each point is from one brain. (C) Left: cartoon to show location of medioposterior cell group in the brain. Right: representative maximum intensity Z-projections of *Phper* and *Phcry2* expression in both hemispheres (L: left, R: right) across CT (individual HCR-FISH channels in rows). Scale bars: 5 µm. (D) Plots of mean transcript abundance, quantified as the number of FISH spots, across CT for *Phper* (left) and *Phcry2* (right). Each point is from one brain. (E, F) As in C and D for anterior-medial a1 cells. Yellow dashed lines: midlines. Scale bars: 10 µm. (G, H) As in C and D for dorsal-lateral cells. Inset text indicates statistical tests (JTK-cycle and RAIN) performed to determine 24-h and 12-h rhythmicity with the tested periods in parentheses. *p* <0.001***, *p* <0.01**, *p* ≤0.05*, *p* >0.05^ns^. For statistics shown in this figure, refer to Supplementary Table 3.

The experiment was repeated with the final tidal agitation occuring slightly later (+1.2 hours) relative to the LD cycle. The animals exhibited robust circatidal activity rhythms, phased to anticipated high water (Figure 5A). Aggregate expression of *per* was again circadian (Supplementary Figure 10A). Across the 10 cell groups examined, the same three showed significant rhythms of one or multiple transcripts (Supplementary Table 4). The expression of *per* and *cry2* in the medioposterior and anterior-medial a1 cells was circadian (Supplementary Figure 10B, C and D, E) whilst the dorsal-lateral cells again showed a strong circatidal, 12-hour, pattern of expression of *per* and *cry*. The transcripts were also oscillating in phase and peaking after subjective high tide (Figure 5B, C). To test whether we could phase-shift these putative tidal cells even further, and to differentiate them from circadian cells exhibiting bimodal 24-hour rhythms, we entrained animals such that the final bout of tidal agitation occurred 5.4 hours phase-shifted relative to the 12:12LD cycle (i.e., subjective high tide at ∼CT3.8 and ∼CT16.2). On release into constant conditions, the animals exhibited appropriately phased circatidal rhythms of activity peaking at subjective high tide (Figure 5A). The aggregate expression of *per* (but not *cry2*) was circadian, peaking in subjective night (Supplementary Figure 11A). Of the 10 cell groups examined, 4 (the original three plus anterior-lateral cells) showed an effect of time on transcript levels (Supplementary Table 5). Medioposterior cells showed antiphasic circadian rhythms of *per* and *cry2*, anterior-medial a1 cells showed a circadian cycle of *per* (Supplementary Figure 11B, C and D, E) and anterior-lateral cells were circadian for *cry2*. In contrast, the dorsal-lateral cells again showed 12-hour cycles of expression of *per* and *cry2* (Figure 5D, E). Moreover, the peaks occurred after subjective high tide and therefore the profiles were phase-shifted relative to the prior LD cycle by ∼6 hours when compared to the previous tidal experiments. Comparison across the three experiments with circatidally active animals showed conservation of the phase relationship between *per* and *cry2* expression as well as adoption of the phase of the prior regime of tidal agitation (Supplementary Figure 12). Thus, in the same brains of *P. hawaiensis* exhibiting free-running circatidal behaviour, medioposterior and anterior-medial a1 cells consistently tracked circadian time, expressed by antiphasic rhythms of *per* and *cry2* expression, whereas the dorsal-lateral cells consistently tracked tidal time, independently of circadian time, and did so with *per* and *cry*2 rhythms in phase.

**Figure 5.**
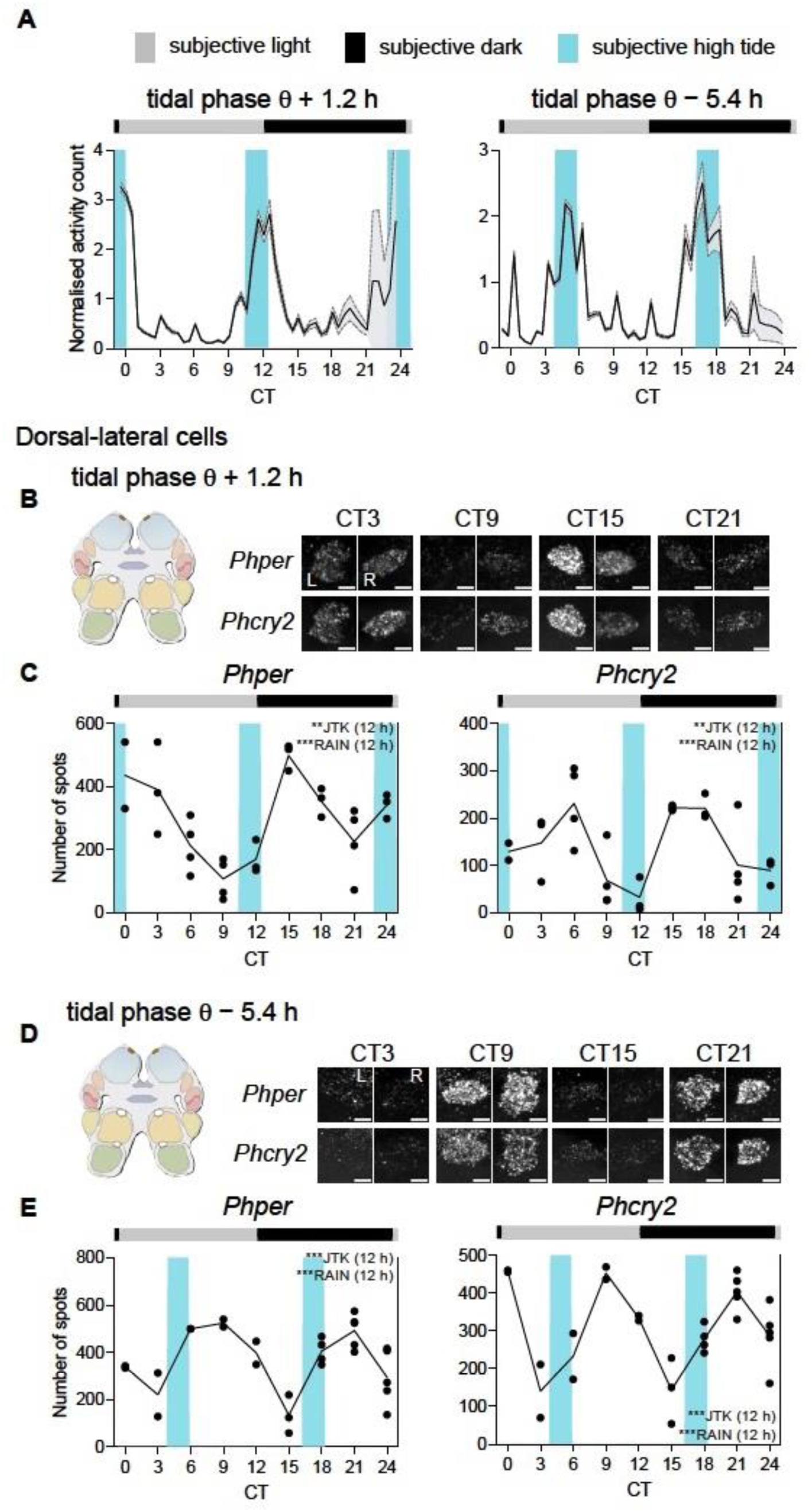
Circatidal rhythms of expression of *Phper*, *Phcry2* in dorsal-lateral cells in the brain of tidally entrained *P. hawaiensis*. (A) Normalised swimming activity (mean +SEM, shading) of *P. hawaiensis* synchronised to the prior agitation cycle, phase-shifted by +1.2 hours (left) or −5.4 hours (right) and harvested for the HCR-FISH time-course (initial group sizes at start of harvest at CT0 = 35). (B) Left: cartoon to show location of dorsal-lateral cell group in the brain. Right: Representative maximum intensity Z-projections of *Phper* and *Phcry2* in both hemispheres (L: left, R: right) across CT (individual HCR-FISH channels in rows). Scale bars: 5 µm. (C) Plots of mean transcript abundance, quantified as the number of FISH spots, across circadian time for *Phper* (left) and *Phcry2* (right) (blue shadings: subjective high water). Each point is from one brain. (D, E) As in B and C for animals entrained to a tidal cycle advanced, relative to the LD cycle, by 5.4 hours. Inset text indicates statistical tests (JTK-cycle and RAIN) performed to determine rhythmicity, with the tested periods in parentheses. *p* <0.001***, *p*< 0.01**, *p* ≤0.05*, *p* >0.05^ns^. For statistics and data of FISH time-course experiments on tidally entrained animals, refer to Supplementary Table 4 and 5 for tidal phases θ +1.2 h and θ –5.4 h, respectively.

### Rhythmic expression of circadian clock genes in daily-rhythmic *P. hawaiensis* under a light:dark cycle

To test whether dorsal-lateral cells have any capacity to follow daily time, we entrained *P. hawaiensis* to LD cycles without tidal agitation. As reported ^13 10^, *P. hawaiensis* exhibited rhythmic activity under LD, peaking at night (Supplementary Figure 13A, B, D, E). Across 7 experiments 58.6 ±4.4% were rhythmic (range: 45 −80%, total N =189). When released into constant darkness, ∼50% of rhythmic animals (i.e., 22.9 ±7.5% of total), showed circadian rhythms in their locomotor activity (Supplementary Figure 13A, C, D, F), with a mean (±SEM) period of 24.17 ±0.17 h (N =39). Activity levels were significantly higher during the subjective night than in the subjective day.

In animals confirmed to be showing daily behavioural activity prior to harvest of their brains for HCR-FISH (Figure 6A), the aggregate expression of *per* was highly rhythmic, peaking at night (Figure 6B). Aggregate expression of *cry2* was also significantly rhythmic, in antiphase to *per*, but with a small amplitude. Of the 10 cell groups examined, 5 showed a daily pattern of one or both transcripts (Supplementary Table 6). In both the medioposterior cell group and the anterior-medial a1 cells, *per* and *cry2* were rhythmic and in antiphase, with nocturnal and diurnal peaks, respectively (Figure 6C, D and E, F), echoing the circadian patterns seen in these cells in tidally active animals under DD. Rhythmic expression of *per* was also evident in the dorsal cells and the anterior-medial b cells (Supplementary Figure 14A, B and C, D), again peaking in the night, and *cry2* was rhythmic in anterior-medial c cells. In contrast, there was no statistically significant rhythmic expression of either *per* or *cry2* in the dorsal-lateral cells of the daily-behaving animals (Figure 6G, H). Thus, several cell groups can exhibit daily cycles of gene expression in daily-behaving *P. hawaiensis* under LD, including the medioposterior and anterior-medial a1 cells that are also circadian under DD. In contrast, the circatidal gene expression observed in dorsal-lateral cells in tidally rhythmic animals under DD was specific to that condition: in the absence of tidal cues this population of cells did show tidal patterning, nor did they adopt the daily rhythms of expression seen in other cell groups under LD.

**Figure 6.**
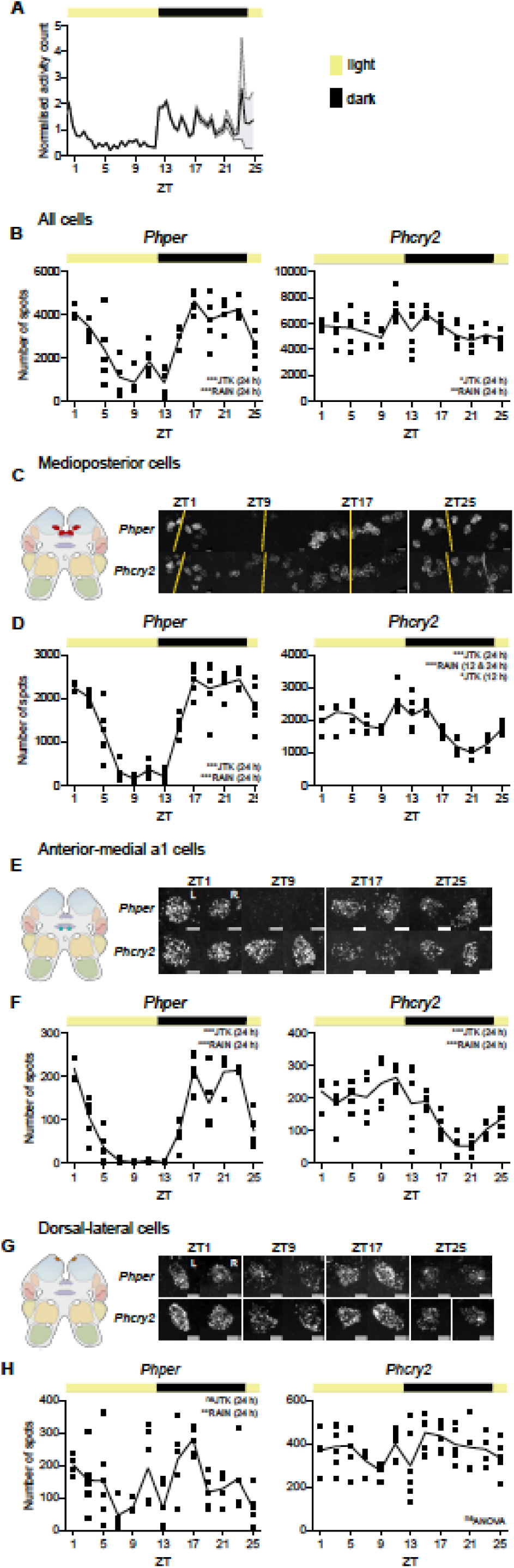
Time course of expression of *per* and *cry2* in cells of *P. hawaiensis* synchronised to the LD cycle. (A) Normalised swimming activity (mean +SEM, shading) of *P. hawaiensis* synchronised to the LD cycle and harvested for the HCR-FISH time-course (initial group size at start of harvest at ZT1 =79). (B) Mean aggregate expression, quantified as the number of FISH spots, of *Phper* and *Phcry2* across all cells plotted across time in animals entrained to, and sampled under, a 12 h:12 h LD cycle. Each point is from one brain. (C) Left: cartoon to show location of medioposterior cell group in the brain. Right: representative maximum intensity Z-projections of *Phper* and *Phcry2* expression in both hemispheres (L: left, R: right) across ZT (individual HCR-FISH channels in rows). Scale bars: 5 µm. (D) Plots of mean transcript abundance, quantified as the number of FISH spots, across daily time for *Phper* (left) and *Phcry2* (right). (E, F) as for C and D for anterior-medial 1a cells. (G, H) as for C and D for dorsal-lateral cells. Inset text in a and c indicate statistical tests (JTK-cycle and RAIN) performed to determine 24-hour rhythmicity. *p* <0.001***, *p* <0.01**, *p* ≤0.05*, *p* >0.05^ns^. See also Supplementary Figure 14 for other cell groups with 24-h cycling in clock gene expression. For statistics and data shown in this figure, refer to Supplementary Table 6.

The distinctions between circadian and circatidal rhythms of gene expression are evident in a combined meta-analysis across the three tidal entrainment studies, registered alternatively in either circadian or circatidal time (Figure 7). When examined across all experiments in the same circatidally behaving animals, the expression of *per* and *cry2* was highly circadian in the medioposterior cells and anterior-medial a1 cells (Figure 7A, B), with *cry2* oscillating in anti-phase to *per*. On the other hand, the expression of *per* and *cry2* in dorsal-lateral cells was highly circatidal and co-phasic (Figure 7C). Circular plots registered to prior LD and tidal agitation emphasise these differences in periodicity and acrophase between the circadian and the circatidal cells (Figure 7D, E). Moreover, this mapping of the tidal oscillations to the phase of tidal entrainment in all three experiments confirmed that the dorsal-lateral cells were able to express a free-running molecular signature of tidal time that was independent of circadian time and revealed, thereby, that the brain of *P. hawaiensis* contains independent circadian and circatidally rhythmic cell populations.

**Figure 7.**
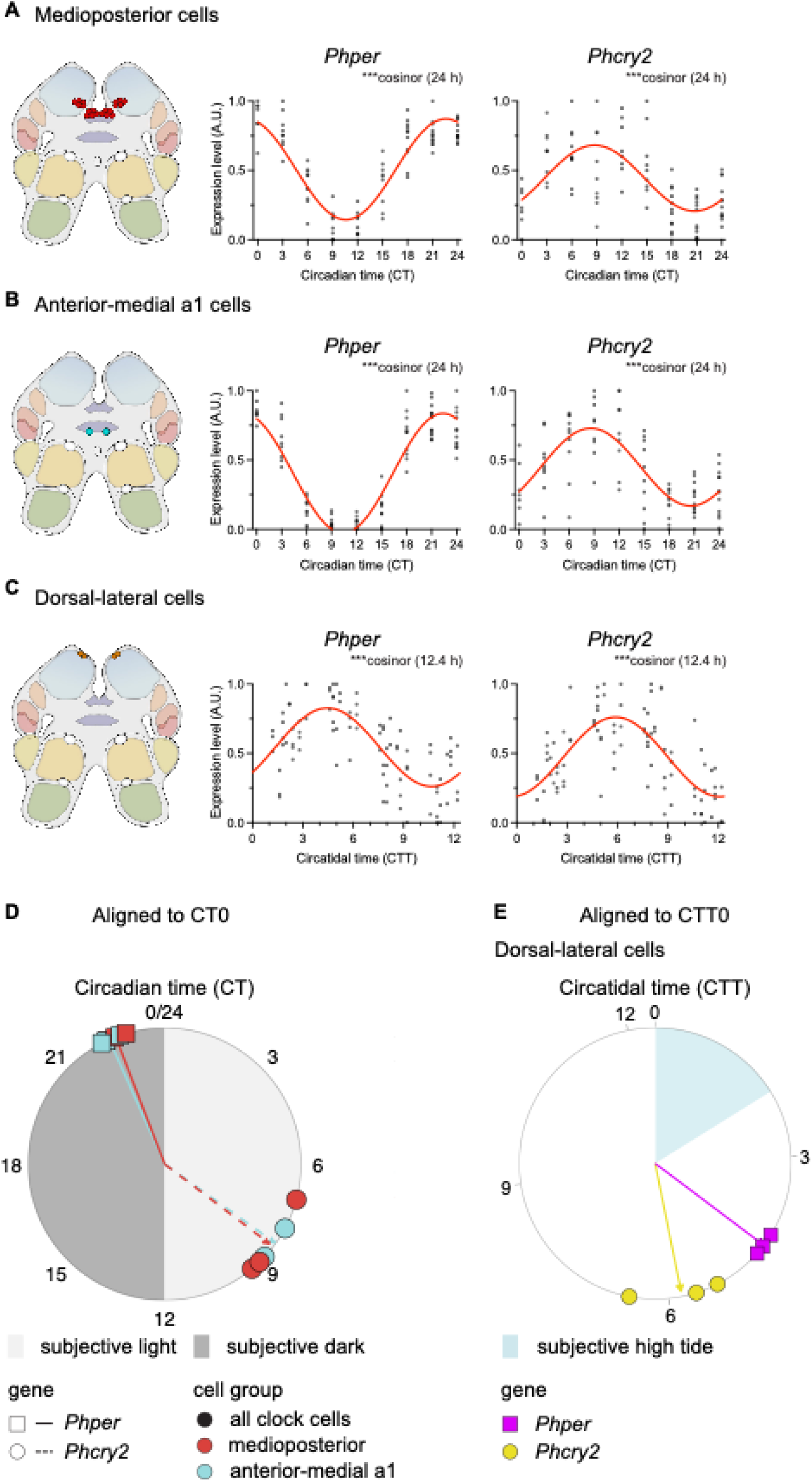
Summary meta-analysis of circatidal and circadian rhythms of clock gene expression in medioposterior, anterior-medial a1 and dorsal-lateral cell groups of *P. hawaiensis*. (A) Left: cartoon to show location of the medioposterior cell group in the brain. Right: scatter plot of expression of *Phper* (left) and *Phcry2* (right) across three independent experiments registered to subjective circadian time (Circadian Time, CT =0). Red lines indicate 24-h cosinor fits. (B) As in A for anterior-medial a 1cell group. (C) As in A for dorsal-lateral cell group but plots are registered to subjective high tide (Circatidal Time, CTT =0). Red lines indicate 12.4-h cosinor fits. Inset text in A-C indicate statistical significance of cosinor (D) Circular plot of circadian peak expression phase of *Phper* and *Phcry2* in selected rhythmic cell groups of *P. hawaiensis* exhibiting circatidal activity rhythms. Each colour-shape combination represents data from across N =3 experiments. Arrowed lines indicate mean phase for each gene-cell group combination, and length of arrowed lines indicates mean resultant length of the vector at each mean. (E) Circular plots of circatidal peak expression phase of *Phper* and *Phcry2* in the dorsal-lateral cells of *P. hawaiensis* exhibiting circatidal activity rhythms, with the circatidal acrophase aligned to CTT0 within each individual experiment. Arrowed lines indicate mean phase for each gene-cell group combination, and length of arrowed lines indicates mean resultant length of the vector at each mean. For mean phase and mean vector lengths for each gene-cell group combination in D and E, see Supplementary Table 7.

## Discussion

Examination of the expression of canonical circadian clock genes across the brains of both *E. pulchra* and *P. hawaiensis* identified ∼60 cells co-expressing *Clk*, *Bmal1*, *cry2* and *per*, arranged into distinct groups across the protocerebrum. Although the loss of *Bmal1* compromises circatidal behaviour in *E. pulchra* ^16^ and in *P. hawaiensis* ^13^, its brain-wide expression showed no local enrichment and so did not highlight any potential circatidal clock cells. This broad distribution is, however, consistent with BMAL1 having diverse, pleiotropic actions beyond timekeeping ^22^. On the other hand, the observed enrichment of *Clk*, *per*, and *cry2* expression (alongside *tim* in *E. pulchra*) in specific cell groups indicated their potential function as circadian oscillators.

In circatidally active *E. pulchra* sampled under LD cycles, the aggregated signals for *per* and *cry2* were daily, reflective of their putative circadian roles, and both peaked in the late night as expected for negative co-regulators within the TTFL. In contrast, aggregate brain expression of *tim* showed significant patterning not only at 24 hours, consistent with circadian cycling of *tim* in whole heads assayed by qPCR ^12^, but also at 12 hours. Analysis at the level of individual cell groups provided a more granular insight, revealing 24-hour rhythms of *per* and *cry2* in medioposterior cells and of *per* in medial cells, with peak expression in late night. This absence of circatidal rhythmicity of *per* expression resonates with the finding that knockdown of *per* to ∼20% of wild-type levels, had no effect on circatidal activity in *E. pulchra*, although it did disrupt circadian phenotypes ^12^. Alternatively, *tim* showed 12-hour accumulation in both the medial and the lateroposterior cells, peaking ∼2 hours after expected high water. These cell-specific differences in *tim* expression likely underlie the compound periodicity evident in the aggregate expression. Furthermore, the 12-hour patterns in the medial and the lateroposterior cells indicate potential tidal time-keeping in these cells associated with circatidal behaviour.

In *P. hawaiensis*, aggregate expression of *per* tracked daily time under LD and circadian time under DD, with elevated levels during the dark phase and subjective night, respectively. Daily and circadian patterning was even more pronounced when examined in individual cell groups, with rhythmic expression of *per* evident in medioposterior, anterior-medial a1, dorsal and anterior-medial b cells. Notably, whereas *per* and *cry2* were in phase in daily cells of *E. pulchra*, expression of *per* and *cry2* in daily and circadian cells in *P. hawaiensis* was antiphasic, suggesting inter-species differences in the underlying molecular architecture of the circadian TTFL.

Remarkably, following entrainment of *P. hawaiensis* to artificial tidal cycles under LD and then subsequent transfer to free-running conditions, dorsal-lateral cells were robustly rhythmic for *per* and *cry2* expression, with a circatidal period that reflected the circatidal behaviour of the animals. Across independent trials, these cells tracked experimental tidal time, regardless of prior LD phase. In contrast, clock gene expression in these cells was arrhythmic when analysed across individuals sampled under LD without tidal cues and with daily, not tidal, behaviour. These observations place the dorsal-lateral cells as strong candidates for dedicated circatidal oscillators, especially when other cell groups in the same brains (e.g., medioposterior and anterior-medial a1 cells), ignored tidal cues and remained circadian, phased to prior LD. Notably, *per* and *cry2* in these tidal cells oscillated in phase, in contrast to the antiphasic rhythms of *per* and *cry2* in daily/ circadian cells. The functional significance of these varying relationships, which are surprising in terms of the canonical TTFL whereby *per* and *cry2* are co-phasic, is not clear but they suggest that a common molecular architecture may have diverged, evolving to operate under different time-bases in different cells. Of the canonical clock genes in *E. pulchra*, only *tim* expression was associated with tidal time in 2 of the 4 neuronal groups studied. For *P hawaiensis* the non-canonical antiphasic expression of *per* and *cry2* occurred in the circadian but not in the circatidal cells. Future analysis of protein expression in these cell groups will better inform understanding of cell-specific TTFL structures. In addition, examination of clock gene expression in the different cell groups of animals with loss/ knock-down of BMAL1 (and CLK?) would indicate mechanistic bridges between the TTFL and tidal behaviour. Nevertheless, it is possible that the circatidal expression patterns of *Phper*, *Phcry2* and *Eptim* are driven by a cryptic tidal clock, such that they reflect cellular activity but are not in themselves causal to tidal behaviour. The genetic tractability of *P. hawaiensis* may enable this to be tested directly.

Notably, even though numerous cells oscillated at ∼24 hours (22 in *P. hawaiensis*) and only a few at ∼12 hours (4 in *P. hawaiensis*), both *E. pulchra* and tidally entrained *P. hawaiensis* were strongly circatidal in terms of behaviour. This raises the question of why strong tidal phenotypes are reflected by relatively few tidal cells, compared to the strong aggregate cycles of daily/ circadian gene expression. Of course, day/ night cycles are highly predictable in nature, and it is therefore expected that circadian systems will be a prominent and robust feature. Nevertheless, coastal animals have evolved temporally complex behaviours to match their habitats ^11 12 13 10^ cued by multiple stimuli. It is therefore possible that gene expression may be circatidal in additional cell groups in *E. pulchra* and/ or *P. hawaiensis* but it evaded statistical detection in the current study because of the high variance intrinsic to population-based measures. Alternatively, it may be that different tidal cell groups respond to distinct entraining cues and so only a subset of these groups is revealed in any single experimental setting. Examination of gene expression in the brain of *P. hawaiensis* entrained by tidal immersion/ emersion ^13^ with and without tidal agitation may indeed reveal additional putative circatidal clock cells: can subsets of cells we describe here as non-rhythmic be recruited by tidal stimuli of other modalities. Alternatively, it may emphasise the distinct and unique tidal role of dorsal-lateral cells. Such a cell- and stimulus-specific entrainment repertoire raises the question of how tidal stimuli are transduced to the central brain and whether a distributed network broadens the capacity for tidal entrainment, reinforcing it in diverse natural settings ^2 10 11^.

In considering the three hypotheses of circatidal time-keeping ^4 5 6^, our data concur most with the model of dedicated tidal and daily timers proposed by Naylor ^6^, with canonical circadian genes in distinct cell groups tuning to either circatidal or circadian periodicities. Our demonstration that dual circatidal and circadian systems operate in parallel also corroborates other work on circatidal clocks. For example, surgical removal of optic lobes of the mangrove cricket, *Apteronemobius asahinai*, abrogates circadian but not circatidal locomotor phenotypes, indicative of two anatomically separable timers ^23 24^. In contrast, evidence in favour of a circadian/ tidal timer based on a single “plastic” clock, as proposed by Enright ^4^, received support from studies on the coastal oyster *Crassostrea gigas*, which showed ∼12-hour and/ or ∼24-hour expression of circadian gene homologues ^25^. Importantly, these models are not mutually exclusive. Given the evolutionary distance between molluscs and arthropods, it is entirely possible that tidal time-keeping machineries have evolved independently. In fact, a parsimonious explanation of the evolution of clock cells with different periodicities would be that pre-existing circadian clock components were co-opted to track relevant tidal cues. The cells we identified invariably co-expressed the rudiments of a circadian TTFL but with either daily or tidal frequencies in the same brain and it is possible that they evolved in parallel, diverging under the influence of photic and mechanosenory (tidal) inputs, respectively.

Kwiatkowski et al. ^10^ asked how *P. hawaiensis* ‘chooses’ between 12.4-hour and 24-hour environmental rhythms during entrainment. Our observation that distinct circadian and circatidal cells groups entrain independently to light and mechanical stimuli, respectively, provides a solution to this problem. Similarly, the authors also questioned whether neurons coordinating rhythmic behaviour in *P. hawaiensis* have two different timers with differential sensitivity to light and tides, or two cell types exist. Whilst we cannot currently exclude the former, our data align better with the latter. This does not preclude interactions of the two systems, however, and the genetic and behavioural malleability of *P. hawaiensis* should enable their decipherment. Thus, our findings may facilitate the examination of input and output pathways of tidal and daily cues at cellular resolution, progressing towards defining the nature and connectivity of circadian and circatidal neural clockworks.

## Acknowledgements

We thank Dr. Sebastian Cachero (LMB) and Dr. Gregory Jefferis (LMB) for their guidance on reference brain generation and their algorithms, as well as Dr. Aziz Aboobaker (Oxford) for providing a starting culture of Chicago-F *P. hawaiensis* and Dr. Michalis Averoff (Lyon) for advice on working with *P. hawaiensis*. We also thank the Mechanical and Electronics Workshops and the Light Microscopy Facility at the LMB for technical support. This work was supported by core funding from Medical Research Council (MRC), as part of United Kingdom Research and Innovation (also known as UK Research and Innovation) (MRC File Reference No. MC_U105170643) to M.H.H. C.Y.S. was supported by a Boehringer Ingelheim Fonds PhD fellowship (2023-2025). For the purpose of open access, the author has applied a CC BY public copyright licence to any Author Accepted Manuscript version arising.

## Author contributions

All Authors contributed to the direction of the project and the writing of the manuscript.

AO and CYS designed and conducted behavioural and neuroanatomical experiments and analysed data.

DCW conducted neuroanatomical experiments and analysed data.

MHH arranged funding.

## Methods

### *Eurydice pulchra* collection and phenotypic analysis

Animals were collected from Llanddona beach, Anglesey, UK, during spring high tides during the months May to October ^12^. Swimming behaviour of both females and males was recorded in DAM10 and LAM10 activity monitors (Trikinetics Inc., USA) ambient lighting (LD) at 16°C with animals housed individually in 3 ml (Ø: 11 mm) plastic vials containing 33 psu artificial seawater (ASW, Reef Salt, Tropical Marine Centre, Germany) and ∼5 mm depth of fine sand. Infrared beam interruptions were recorded using the DAMSystem3 software (TriKinetics Inc., USA) in one-minute bins. The first peak in tidal swimming activity was omitted from analysis and behavioural rhythmicity was assessed over the subsequent six tidal cycles using chi-squared periodogram analysis in Rethomics R package ^26^.

### *Parhyale hawaiensis* husbandry and behavioural analysis

Chicago F strain *P. hawaiensis* were maintained in 6.5 L polypropylene containers (h: 125 mm, w: 260 mm, d: 260 mm) containing 30 –35 psu ASW and a layer of crushed coral substrate (grain size 2 –20 mm). Animals were reared at 25°C under 12h light:dark (LD) cycles. The containers were covered with clear Perspex lids to prevent evaporation and aerated using aquarium air pumps and air stones. Animals were fed carrots, goldfish flakes (Tetra GmbH, Germany), algae wafers (Kyorin Food Ind. Ltd., Japan), freeze-dried brine shrimps, and/or freeze-dried Tubifex (Interpret, UK) once weekly. Full water changes were performed fortnightly. For locomotor activity monitoring, female and male animals were placed individually into 23 mL polystyrene vials (h: 75 mm, Ø: 23.5 mm) containing a layer ( ∼2 mm) of crushed coral substrate (grain size 2 –5 mm) and approximately 6 cm of 30 psu ASW at 25°C. Tubes were loosely covered with parafilm and loaded into two-tier activity monitors modified from the LAM25 Locomotor Activity Monitors (TriKinetics Inc., USA; Supplementary Figure 2a) configured to record swimming and/or roaming activity. Infrared beam interruptions were recorded using the DAMSystem3 software (TriKinetics Inc., USA).

Beam interruptions were collected into 30-minute bins and normalised within each animal by dividing the count in each bin by the average count across all bins for that animal. Autocorrelation periodogram was used to determine whether animals were rhythmic, with a significance threshold of *α* = 0.05 ^27^. The period and the power of rhythmicity were determined using Lomb-Scargle periodogram analysis ^28^ of binned and normalised data at the significance level *α* =0.05. For circatidal experiments, the first subjective high tide after loading was omitted from analysis and behavioural rhythmicity was assessed over the subsequent six tidal cycles.

For circadian entrainment experiments, animals loaded into the monitor were subjected to at least four 12:12 LD (ZT0/lights-on at 0600 local time) cycles before being allowed to free run in DD. For tidal entrainment, animals were kept under LD as above but mechanically agitated on an orbital shaker set at 50 –60 rpm for 2 hours at 12.4-hour intervals. For behavioural monitoring, animals were taken from entrainment regimes during the agitation phase and placed in the activity monitor as described above to free-run without agitation in DD.

### Brain collection and processing for immunohistochemistry and HCR-FISH

Animals were anaesthetised in ice-cold seawater, and their brains dissected using sharpened watch-maker’s forceps in ice-chilled physiological saline ^29^ (*E. pulchra)* or 0.22 µm filter-sterilised ASW (30 −35 psu, FASW; *P. hawaiensis*). Tissues were immediately fixed in 4% paraformaldehyde (made in PBS for *E. pulchra* and FASW for *P. hawaiensis*) for 1 hour at room temperature, dehydrated serially in 33%, 66%, 100% methanol in PBST (PBS supplemented with 0.1% Tween20), and stored in 100% methanol at −20°C for at least 16 h before hybridisation or immunohistochemistry. Immediately prior to use, brains were rehydrated in 66/33% methanol and washed/equilibrated in PBST (3×10 min). Neuropils were immunolabelled using anti-SYNORF1 antibody (3C11, Developmental Studies Hybridoma Bank (DSHB)). For *E. pulchra* and *P. hawaiensis* brains immunohistochemistry followed ^30, 31^ and (Supplementary Methods). HCR^TM^RNA-FISH probes were designed and synthesised by Molecular Instruments Inc, USA against user-supplied sequences (Supplementary Table 8). HCR-FISH followed the manufacturer’s protocol ^32^ with some minor alterations following ^33^ (Supplementary Methods). To co-localise putative clock cells relative to major neuropils, immunostaining with anti-SYNORF1 antibody was followed by HCR-FISH (Supplementary Methods). Reference brains were generated from confocal image stacks of brains in which neuropils were labelled with anti-SYNORF1 (Supplementary Methods).

### Time-course sampling design

To investigate clock gene expression pattern in the brain of *E. pulchra*, animals were collected from Llanddona beach, Anglesey, UK, during spring high tides in October 2024 and loaded into activity monitors. Prior to sampling, animals were maintained under 18 h: 6 h L:D cycle for 24 hours. Animals showing peaks of swimming activity during the two subjective high waters during this interval were sampled every two hours, beginning at ZT7 (Supplementary Figure 15A). To investigate circatidal clock gene expression patterns in the brain of tidally entrained *P. hawaiensis*, animals which had been subjected to tidal entrainment were loaded into activity monitors as described. Sampling occurred every 3 hours, beginning at CT0 on the day after monitor loading and ending at CT24, for a total of 9 timepoints (Supplementary Figure 15B). Individuals were only sampled if they showed peaks of locomotor activity during the subjective high tides preceding their sampling timepoint. To investigate daily clock gene expression pattern in the brain of *P. hawaiensis*, animals were loaded into activity monitors and subjected to 12 h: 12 h LD cycle for 6 days as described. Individuals showing significant 24-h rhythmicity in locomotor activity (as described) over this entrainment period up to the day before sampling were marked for sampling. On the day of sampling, marked animals were sampled every 2 hours, beginning at ZT1 and ending at ZT25, for a total of 13 timepoints (Supplementary Figure 15C). Dissected brains were processed for digital HCR-FISH as described. HCR-FISH probes used for time-course experiments are described in Supplementary Table 10.

### Whole brain digital HCR-FISH

For quantification of diffraction-limited spots ^32^, signal amplification with 60 nM of each hairpin was performed for 1 h, followed by staining with 1 µg/mL DAPI (as above). Brains were mounted in Prolong Glass Antifade Mountant (ThermoFisher #P36980). *E. pulchra* brains were mounted with the posterior surface apposed to the coverslip whilst *P. hawaiensis* brains were mounted with the anterior surface apposed. To minimise signal loss with imaging depth, brains were compressed to within 40 µm by mounting the coverslips containing brains onto spacers made from double-sided adhesive tape (Radio Spares #770-3422). An Andor BC43 CF benchtop microscope equipped with a 60x/1.42NA Plan Apochromat oil immersion objective was used to image each brain with an optical section interval of 0.3 µm.

To prepare images for segmentation of cell groups enriched for expression of *per*, *cry2*, and/ or *Clk/ tim*, 2D maximum filters were applied to each HCR-FISH channel, followed by summation of the filtered images across channels and 2D gaussian filtering. The resulting images, together with the corresponding raw DAPI image as the auxiliary channel, were subjected to segmentation using Cellpose ^34^ and a custom model (available at GitHub) trained via the human-in-the-loop functionality of the Cellpose 2.0 graphical user interface ^35^. The resulting masks were manually curated as required in napari ^36^ and assigned to distinct cell groups. For spot detection, images were deconvolved using ClearView^TM^ Deconvolution with 10 iterations after being acquired in the Fusion software. RNA foci were then detected in deconvolved images and assigned to cell groups using the RS-FISH ImageJ plugin ^37^.

### Statistical analyses

Shapiro-Wilk test used to determine normal distribution of data, and Levene’s test used to determine if the variances of all groups are equal. An overall time effect was tested by ANOVA for normally distributed data and by Kruskal-Wallis for non-normal datasets. Where this yielded a statistically significant effect, rhythmicity in the data was analysed further by JTK CYCLE ^20^ and RAIN ^21^ at 12-hour or 24-hour periods. JTK-cycle p-values are adjusted using Bonferroni’s method, and RAIN p-values are adjusted using the adaptive Benjamini-Hochberg (default) method. Gene expression was considered rhythmic when ANOVA was significant, and both JTK CYCLE and RAIN were significant for the same period (12 hours or 24 hours).

## Supplementary materials

### Supplementary Figures and Legends

**Supplementary Figure 1.**
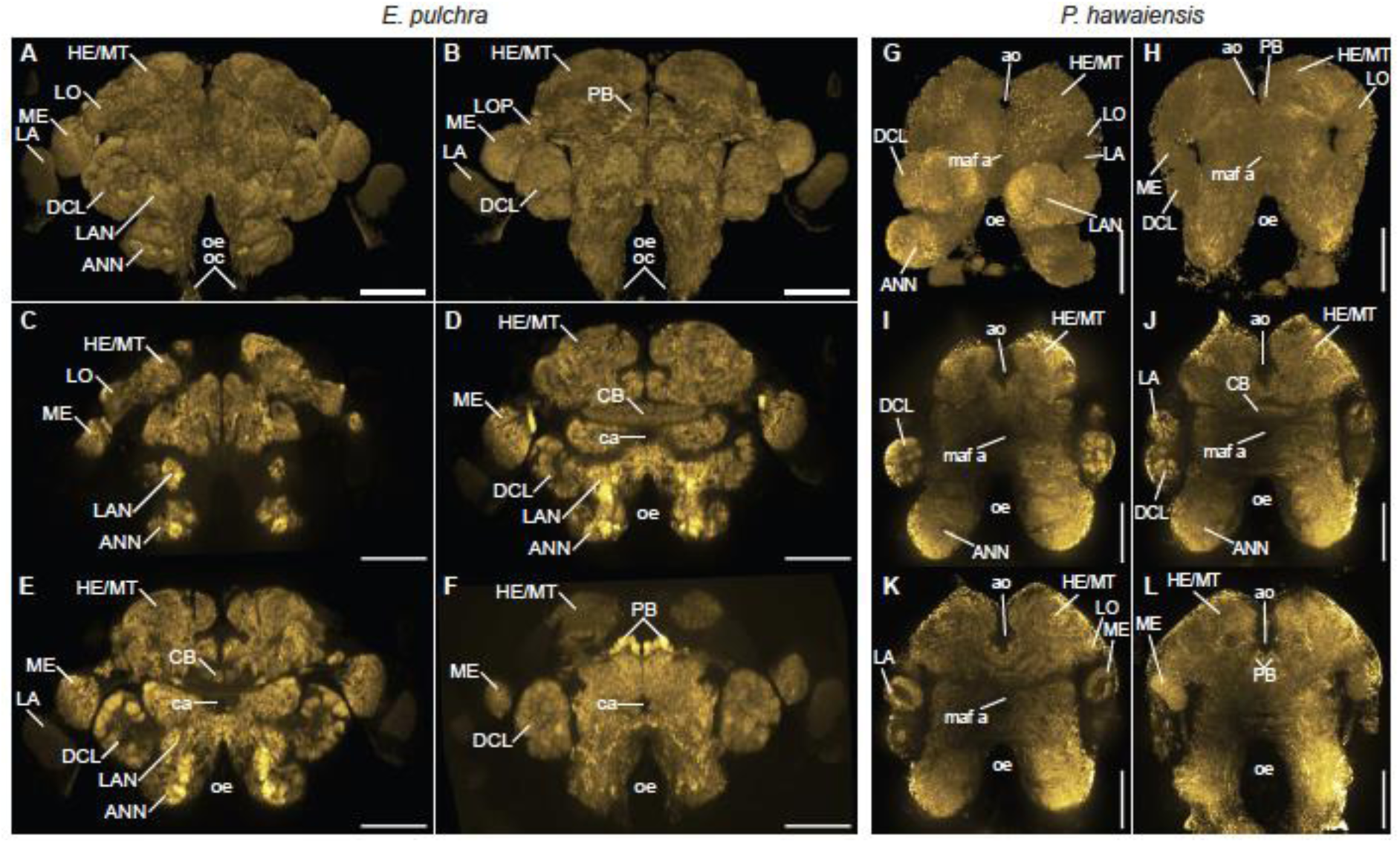
Overview of representative *E. pulchra* and *P. hawaiensis* brains, immunostained with anti-SYNORF1 to highlight the neuropils. (A) Anterior view of a *E. pulchra* brain (3D volume projection), with optic lobes, deutocerebral chemosensory lobes, antenna neuropils and hemi-ellipsoid body/medulla terminalis complexes are visible on the anterior surface of the brain. (B) Posterior view of a *E. pulchra* brain (3D volume projection). The optic lobes, deutocerebral chemosensory lobes and hemi-ellipsoid body/medulla terminalis complexes are visible on the posterior surface. The protocerebral bridge appears as a distinct arch-like structure under the hemi-ellipsoid body/medulla terminalis complexes. (C-E) Selected optical sections of a *E. pulchra* brain from anterior to posterior, showing locations of distinct neuropils throughout the depth of the brain. The optic lobes span the anterior-posterior axis, with the lobula constrained to the anterior aspect (C), and the medulla and lamina (D, E) occupying the medial and posterior aspects. The cigar-shaped central body is at its most expansive towards the middle of the brain. (F) Maximum intensity Z-projection through the protocerebral bridge of a *E. pulchra* brain. (G) Anterior view of a *P. hawaiensis* brain (3D volume projection), with parts of the optic lobes (lamina and lobula), deutocerebral chemosensory lobes, antenna neuropils and hemi-ellipsoid body/medulla terminalis complexes are visible on the anterior surface of the brain. (H) Posterior view of a *P. hawaiensis* brain (3D volume projection). Parts of the optic lobes (medulla and lobula), deutocerebral chemosensory lobes and hemi-ellipsoid body/medulla terminalis complexes are visible on the posterior surface. As in *E. pulchra,* the protocerebral bridge of *P. hawaiensis* is located posteriorly under the hemi-ellipsoid body/medulla terminalis complexes. (I-K) Selected optical sections of a *P. hawaiensis* brain from anterior to posterior, showing locations of distinct neuropils throughout the depth of the brain. The hemi-ellipsoid body/medulla terminalis complexes and the foramina accommodating two of the brain arteries, the anterior aorta and myoarterial formation a, are visible throughout the thickness of the brain. The former also constitute the most dorsal neuropils of the brain. The deutocerebral chemosensory lobes are located anterior to the rest of the optic lobes (I, J). The lamina is the most anteriorly positioned optic neuropils (J), followed almost simultaneously by the medulla and lobula (K). The sausage-shaped central body is at its most expansive towards the middle of the brain (J). (l) Maximum intensity Z-projection through the protocerebral bridge of a *P. hawaiensis* brain. *Abbreviations*: ANN, antenna 2 neuropil; ao, anterior aorta; ca, cerebral artery; CB, central body; DCL, deutocerebral chemosensory lobe; HE/MT, hemi-ellipsoid body/medulla terminalis; LA, lamina; LAN, lateral antenna 1 neuropil; LO, lobula; LOP, lobula plate; maf a, myoarterial formation a; ME, medulla; oc, oesophageal connective; oe, oesophageal foramen; PB, protocerebral bridge. Scale bars: 100 μm.

**Supplementary Figure 2.**
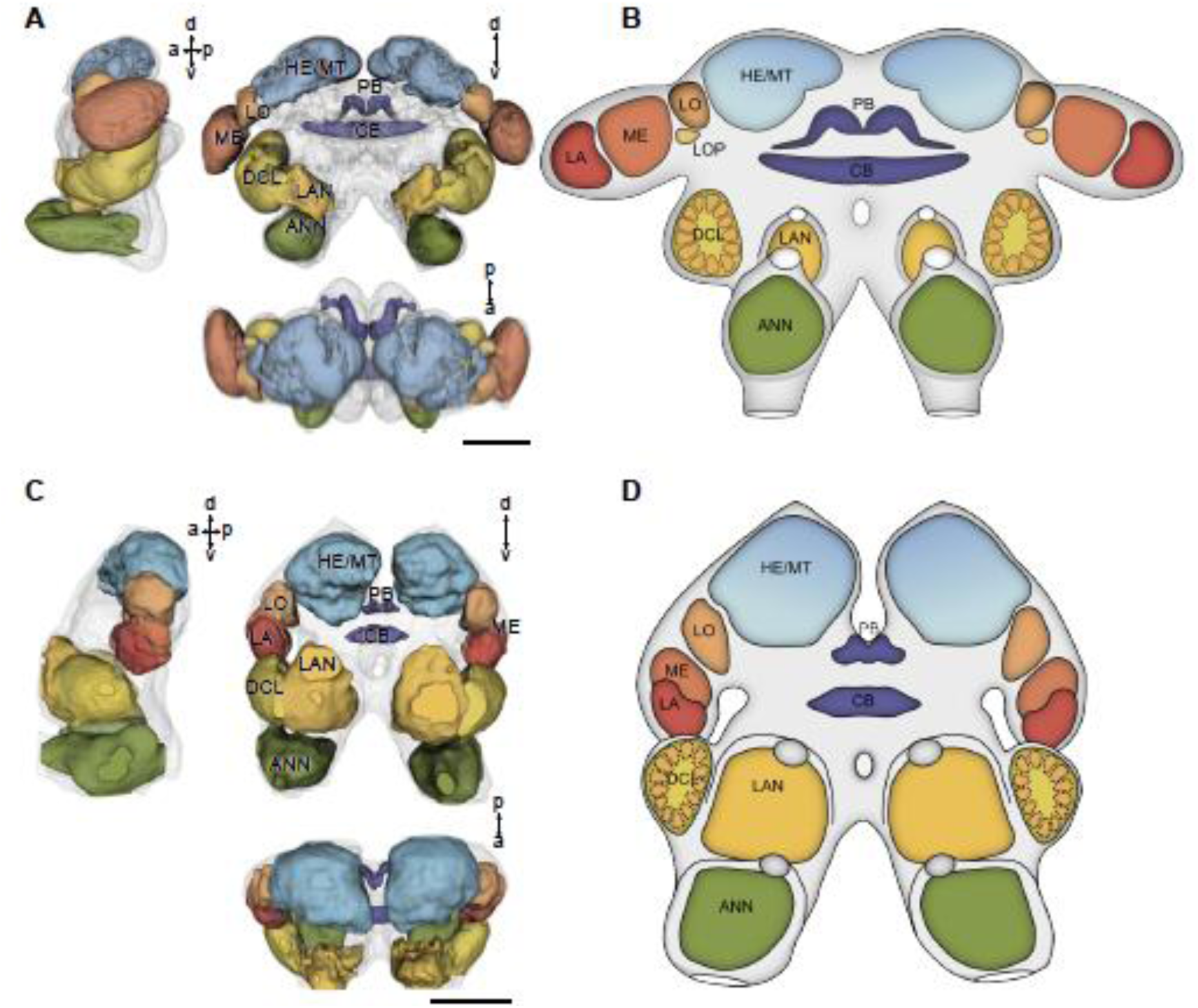
Gross anatomy of *E. pulchra* and *P. hawaiensis* brains revealed by anti-SYNORF1 immunostaining, registered to refence brain. (A) Lateral, anterior and dorsal views of the *E. pulchra* surface-rendered 3D average reference brain, with major neuropils demarcated. See also Supplementary Video 1. (B) Diagrammatic representation of the of *E. pulchra* brain, based on A. (C) Lateral, anterior and dorsal views of *P. hawaiensis* surface-rendered 3D average reference brain with major neuropils demarcated. See also Supplementary Video 2. (D) Diagrammatic representation of the *P. hawaiensis* brain, based on C. Compass markers in A and C show anterior (a), posterior (p), dorsal (d) and ventral (v) directions. *Abbreviations:* ANN antenna 2 neuropil, CB central body, DCL deutocerebral chemosensory lobe, HE/MT medulla terminalis/ hemi-ellipsoid body, LA lamina, LAL lateral accessory lobe, LAN lateral antenna 1 neuropil, LO lobula, LOP lobula plate, ME medulla, PB protocerebral bridge. Scale bars: 100 µm.

**Supplementary Figure 3.**
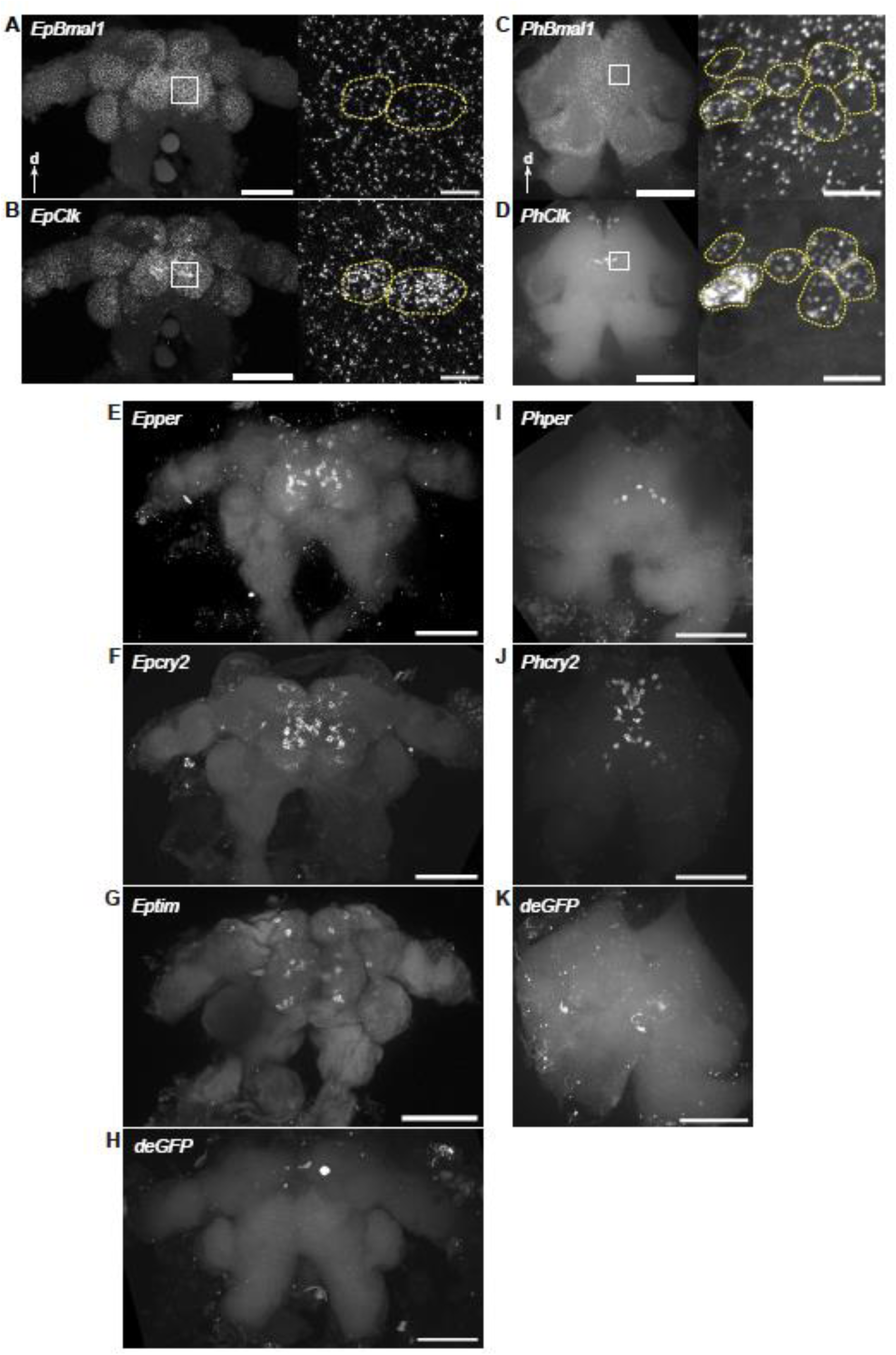
Expression of circadian clock positive and negative regulators across the brains of *E. pulchra* and *P. hawaiensis*. (A) Left panel: maximum intensity Z-projection of image stack of an entire *E. pulchra* brain (posterior apposing coverslip) probed for circadian factor *EpBmal1* using HCR-FISH. Right panel: maximum intensity Z-projection of two-cell-thick image stack (16 µm) from the boxed region of the brain in the left panel. (B) As in A, probed for circadian factor, *EpClk*. (C) Left panel: maximum intensity Z-projection of image stacks of a *P. hawaiensis* brain (anterior apposing coverslip) probed for circadian factor *PhBmal1*. Right panel: maximum intensity Z-projection of two-cell-thick image stacks (24 µm) from the boxed region of the brain in the left panels. (D) As in C, probed for circadian factor, *PhClk*. (E, F, G, H) Maximum intensity Z-projections of image stacks of *E. pulchra* brains (posterior apposing coverslip) probed for circadian clock repressors, *Epcry2, Epper*, *Eptim* and the negative control probe *deGFP* HCR-FISH. (I, J, K) Maximum intensity Z-projections of image stacks of *P. hawaiensis* brains (anterior apposing coverslip) probed for circadian clock repressors, *Phcry2*, *Phper* and the negative control probe *deGFP*. Scale bars: 100 µm. Arrows indicate dorsal orientation. Yellow dotted outlines in A-D indicate cells in which *Clk* expression is enriched. Scale bars: 100 µm (A-D: left panels, E-K) and 10 µm (A-D: right panels).

**Supplementary Figure 4.**
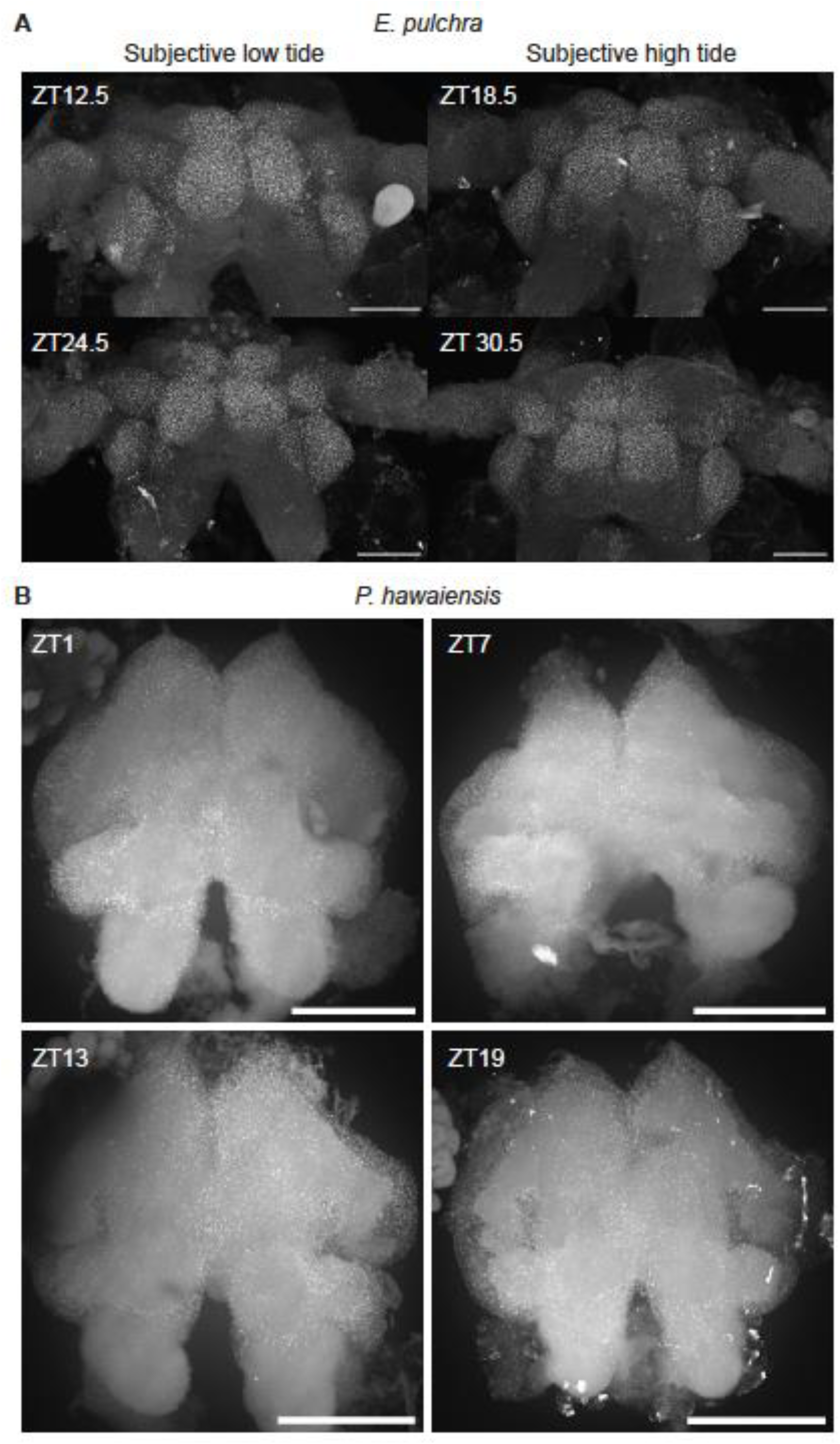
Cellular expression of *Bmal1* in brains of *E. pulchra* and *P. hawaiensis* across time. (A) Maximum intensity Z-projections of representative brains (posterior apposing coverslip) from field-collected *E. pulchra* sampled under 14 h: 10 h LD cycle and then probed for *EpBmal1* using HCR-FISH. (B) Maximum intensity Z-projections of representative *P. hawaiensis* brains (anterior apposing coverslip) sampled under 12 h: 12 h LD cycle and then probed for *PhBmal1* using HCR-FISH. Scale bars: 100 µm.

**Supplementary Figure 5.**
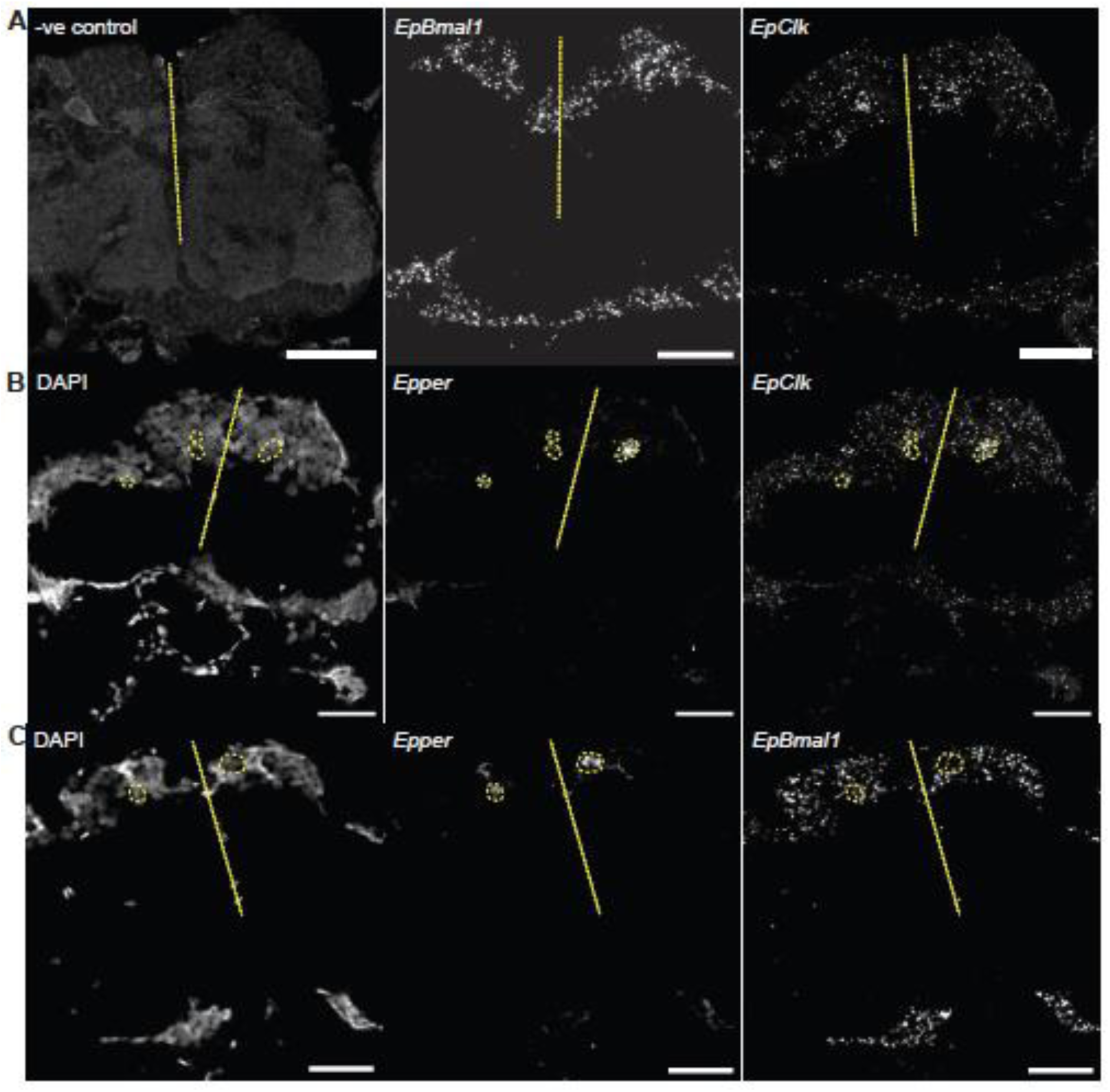
Expression of circadian clock genes in *E. pulchra* brain sections detected using RNAScope^TM^. (A) Representative image (single optical section) of *E. pulchra* brain sections probed for negative control (left), *EpBmal1* (middle) and *EpClk* (right). (B) Representative image (single optical section) of a *E. pulchra* brain section stained with DAPI (left) and probed for *Epper* (middle) and *EpClk* (right). (C) As in B for *Epper* and *EpBmal1*. Yellow dashed outlines indicate cells enriched for *Epper* and *EpClk* expression. Yellow dotted lines indicate midlines. Scale bars: 10 µm.

**Supplementary Figure 6.**
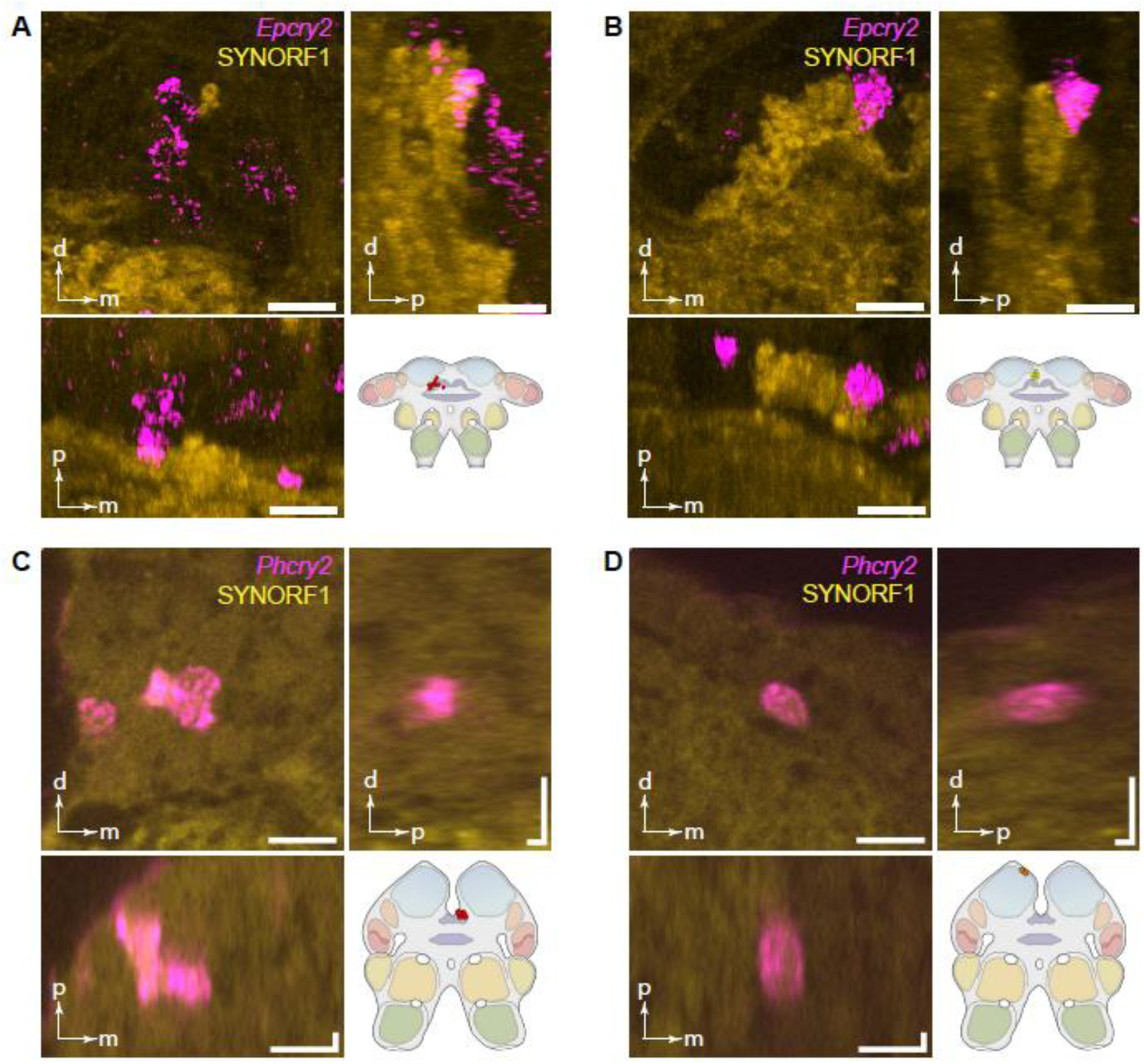
Representative orthogonal views of brains co-labelled with anti-SYNORF1 (IHC) and *cry2* (HCR-FISH). (A) Orthogonal views showing *Phcry2* enrichment in four medioposterior cells of a *E. pulchra* brain co-stained with anti-SYNORF1. Cartoon shows location of these cells. (B) As in A but for the medial cells. (C) Orthogonal views showing *Phcry2* enrichment in four medioposterior cells of a *P. hawaiensis* brain co-stained with anti-SYNORF1. Cartoon shows location of these cells. (D) As in C but for the two dorsal-lateral cells. Horizonal (and vertical) scale bars: 10 µm. Orientations: d: dorsal; p: posterior; m: medial.

**Supplementary Figure 7.**
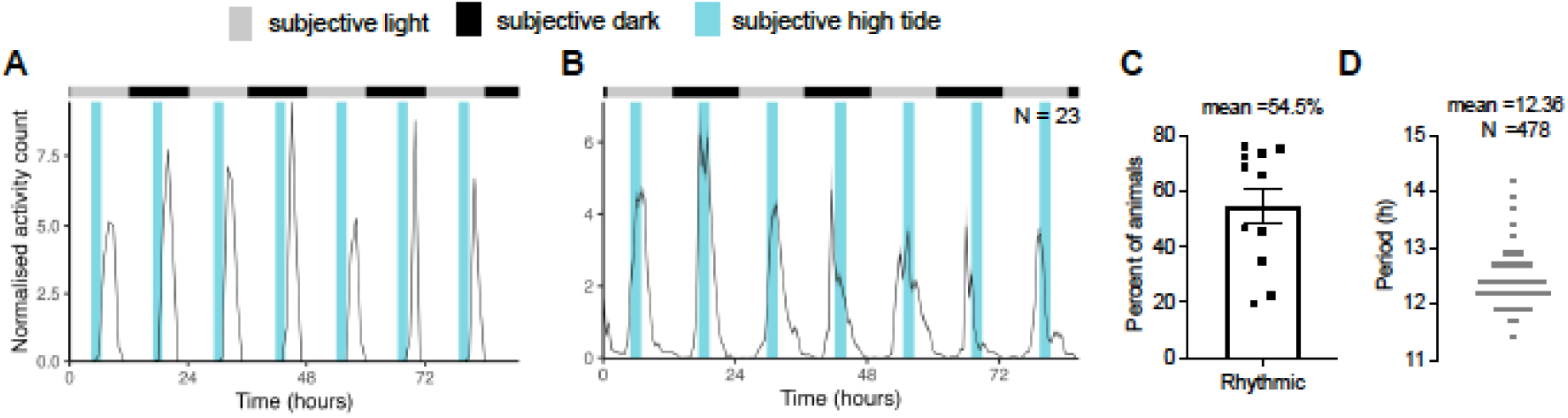
Circatidal swimming behaviour of beach-collected *Eurydice pulchra* recorded in free-running laboratory conditions. (A) Normalised swimming activity of a representative individual *E. pulchra*. Blue shadings indicate subjective high tides on home beach. (B) Mean normalised swimming activity profile for a group of animals under DD from the same experiment as the individual in A (N =23). (C) Percentage of animals showing rhythmic swimming behaviour from 11 independent experiments, representing 11 beach collections from 2022 to 2024. Error bars indicate SEM. (D) Individual periods of beach-caught *E. pulchra* swimming activity under DD across all experiments. Mean (±SEM) is 12.36 ±0.02 h (N =478).

**Supplementary Figure 8.**
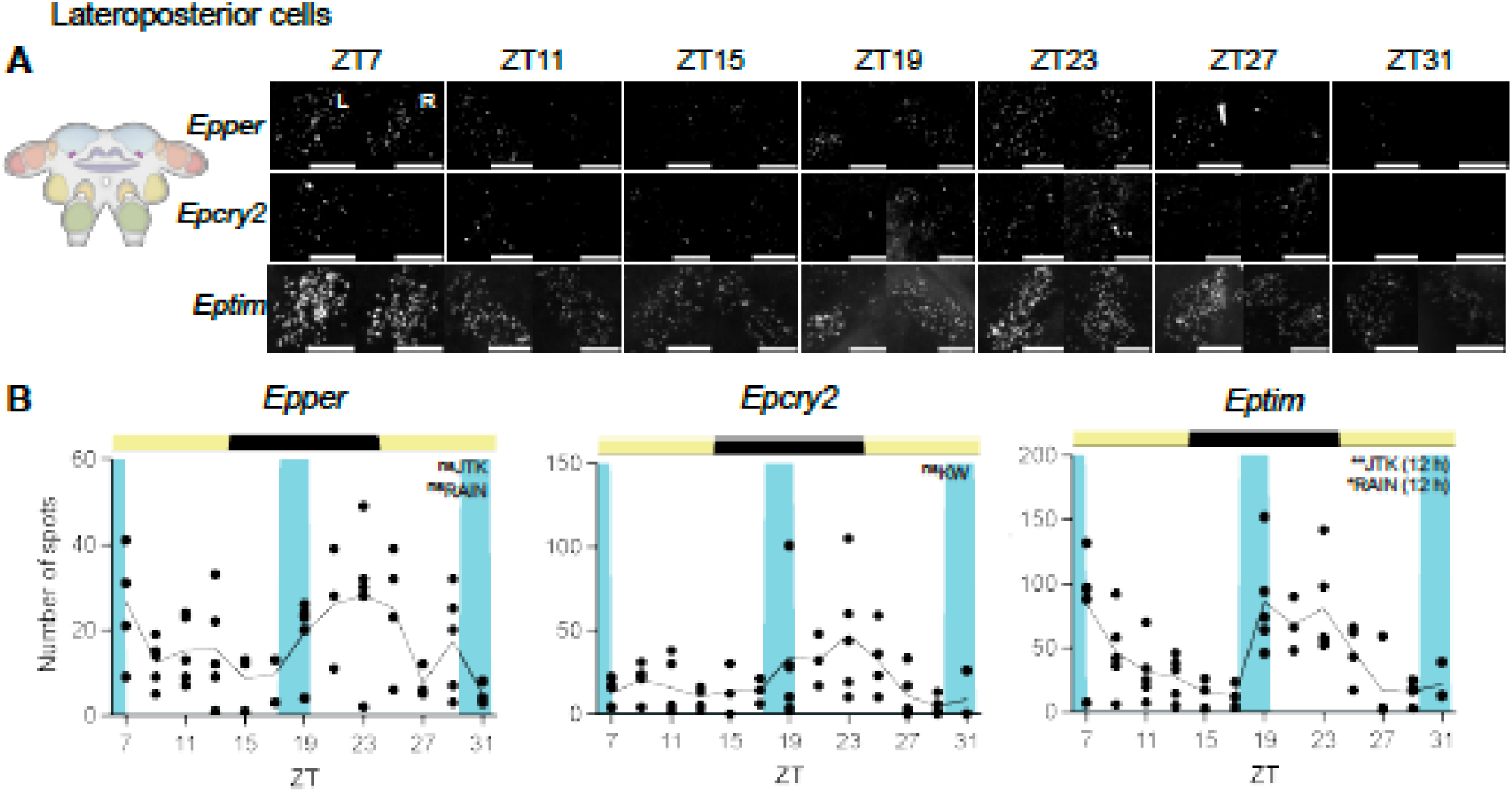
Time course of expression of *per, cry2* and *tim* in lateroposterior cells of tidally rhythmic *E. pulchra* held on a LD cycle. (A) Left: Cartoon to show location of lateroposterior cell group in the brain and Right: representative maximum intensity Z-projections of *Epper*, *Epcry2* and *Eptim* expression in lateroposterior cells in both hemispheres (L: left, R: right) across ZT (individual HCR-FISH channels in rows). (B) Plots of mean transcript abundance, quantified as the number of FISH spots, across ZT for *Epper* (left), *Epcry2* (middle) and *Eptim* n (right) in the lateroposterior cells. Each point is from one brain. Inset text indicates statistical tests performed to determine variation of gene expression with time and/or rhythmicity, with the tested periods (if any) in parentheses. *p* <0.001***, *p* <0.01**, *p* ≤0.05*, *p* >0.05^ns^. For statistics shown in this figure, refer to Supplementary Table 2. Scale bars: 10 µm (A, C).

**Supplementary Figure 9.**
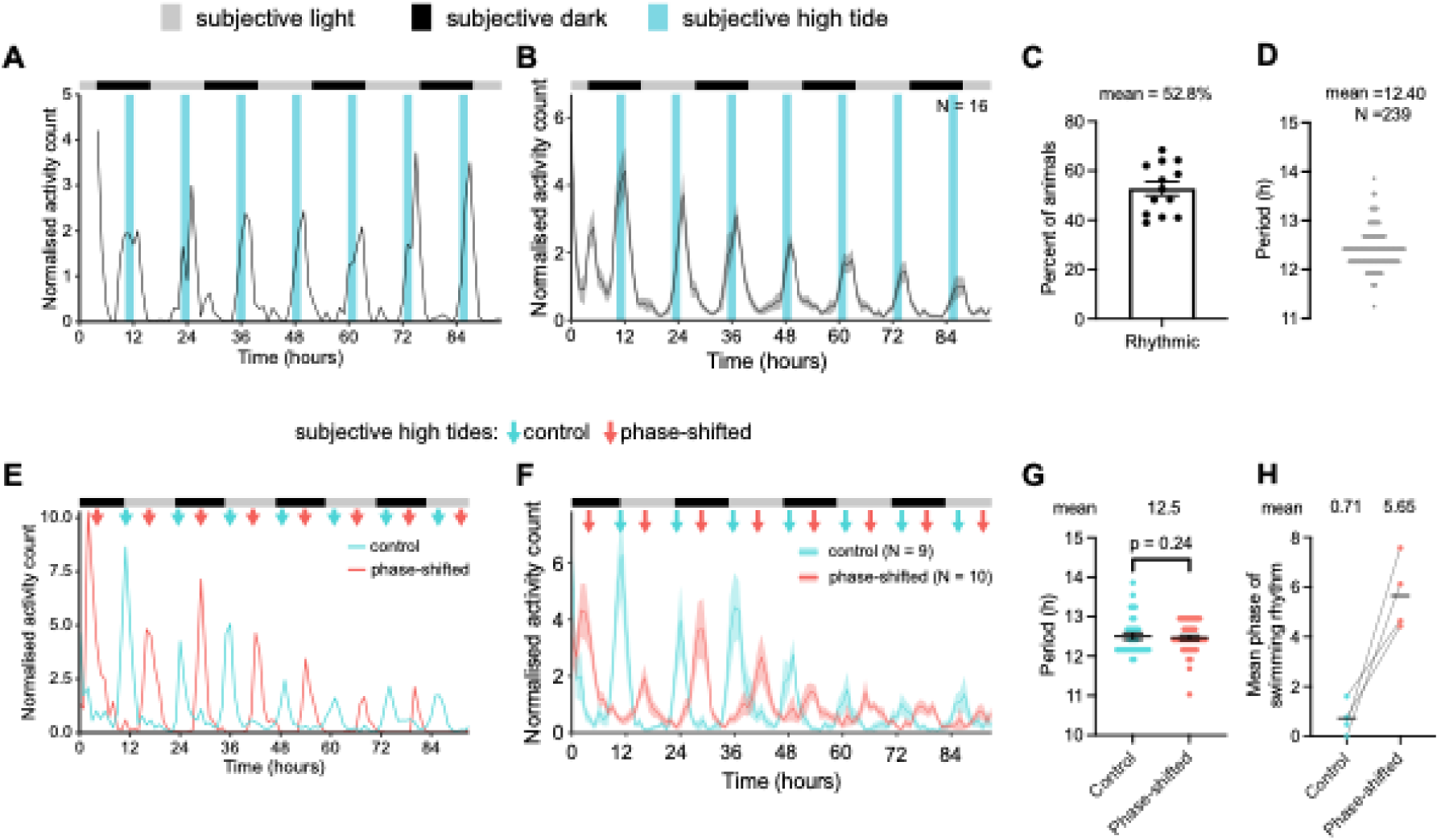
Laboratory-cultured *P. hawaiensis* show circatidal activity rhythms after entrainment to a tidal cycle of 2 hours of mechanical agitation every 12.4 hours simulating high water. (A) Normalised swimming activity of individual laboratory-cultured *P. hawaiensis* free-running under DD (grey/ black bars), following entrainment to 2 h of agitation every 12.4 h. Blue bars indicate time of subjective tidal agitation. (B) Mean normalised swimming activity profile for all circatidal rhythmic *P. hawaiensis* under DD (grey/ black bars) from the same experiment as the individual in A (N =16). Blue bars indicate time of subjective tidal agitation, shaded area indicates ±SEM. (C) Percentage of animals showing a circatidal rhythm in swimming activity after tidal entrainment by agitation. Data from 13 independent experiments (N =456). Error bars indicate SEM. (D) Individual periods of circatidal locomotor activity under DD of tidally entrained *P. hawaiensis.* Mean (±SEM) is 12.40 ±0.02 h (N =239). (E) Representative activity recording of two individual laboratory-cultured *P. hawaiensis* free-running under DD (black/ grey bars) following prior tidal entrainment at a period of 12.4 h (cyan, control) or 12.9 h (red, phase-shifted) over 6 days. Arrows indicate times of subjective tidal agitation for the control and phase-shifted animals. (F) Mean normalised swimming activity profile for all circatidally rhythmic *P. hawaiensis* under DD (grey/ black bars) from the same experiment as the individuals in E (N_control_ =9, N_phase-shifted_ =10). Arrows indicate times of subjective tidal agitation for the control and phase-shifted groups, shaded area indicates ±SEM. (G). Individual periods of circatidal activity rhythm of *P. hawaiensis* after tidal entrainment by agitation under control and phase-shifted conditions (4 independent experiments, N_control_ =34, N_phase-shifted_ =42, Kolmogorov-Smirnov test: *D* =0.24, p =0.24, n.s.). Horizontal lines indicate mean phase within each group, and error bars indicate SEM. (H) Mean phase of circatidal activity rhythms of paired control and phase-shifted groups across 4 replicate experiments as in F. The subjective high tide onset of the earliest control group defined phase =0. Horizontal lines indicate mean phase within each group.

**Supplementary Figure 10.**
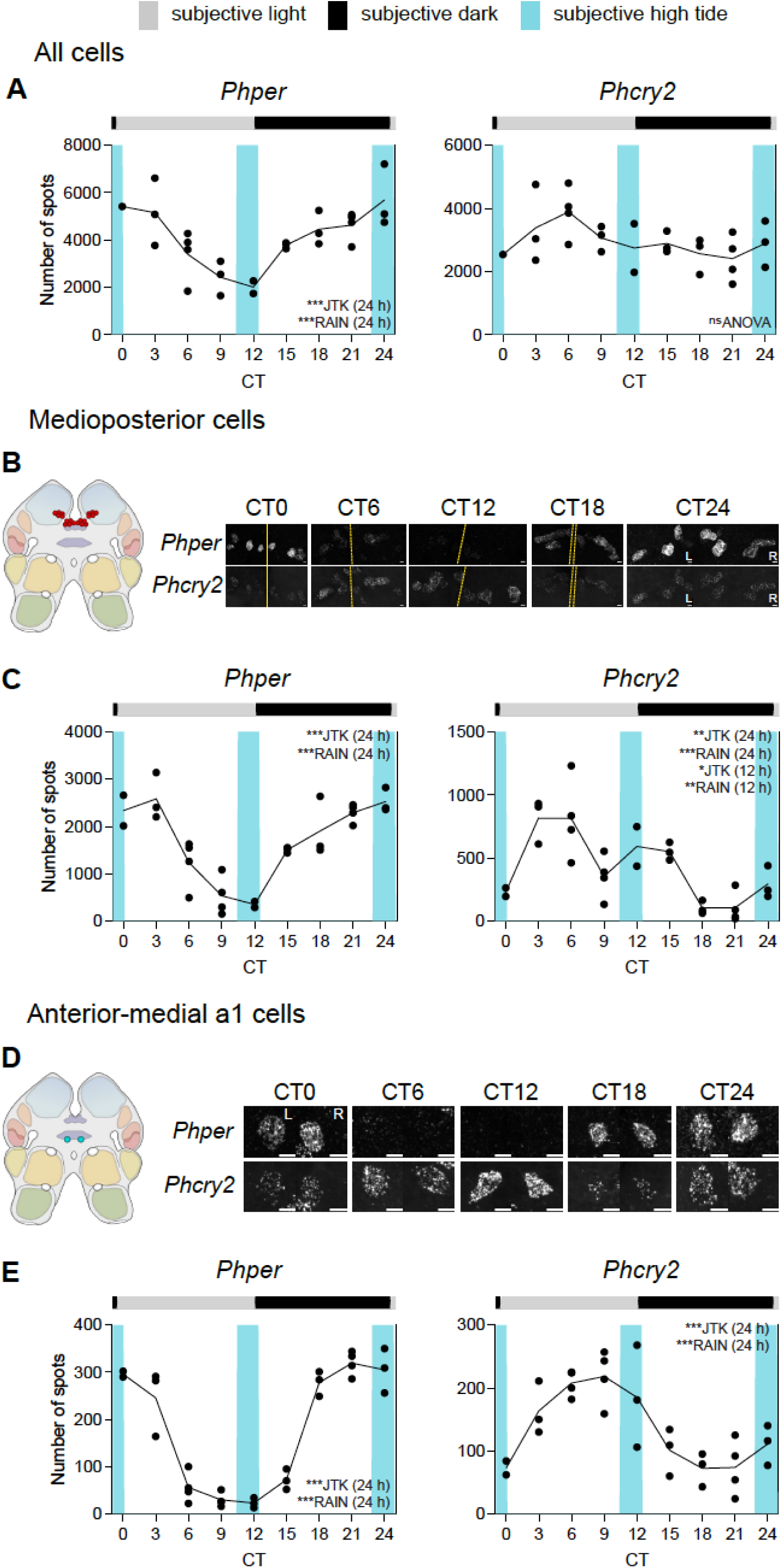
Free-running clock gene expression rhythms in tidally entrained *P. hawaiensis* (tidal phase θ + 1.2 h). (A) Mean aggregate expression, quantified as the number of FISH spots, of Phper and *Phcry2* across all cells plotted across time in tidally entrained animals sampled under DD. Each point is from one brain. (B) Left: Cartoon to show location of medioposterior cell group in the brain and Right: representative maximum intensity Z-projections of *Phper* and *Phcry2* expression in the medioposterior cells from both hemispheres (L: left, R: right, yellow dashed lines: midlines) across CT (individual HCR-FISH channels in rows). (C) Mean transcript abundance, quantified as the number of FISH spots, across CT for *Phper* (left) and *Phcry2* (right)) in the medioposterior cells. Each point is from one brain. (D, E) As in B, C for anterior-medial a1 cell group. Inset text indicates statistical tests performed to determine variation of gene expression with time and/or rhythmicity, with the tested periods (if any) in parentheses. *p* <0.001***, *p* <0.01**, *p* ≤0.05*, *p* >0.05^ns^. KW: Kruskal-Wallis test. For statistics shown in this figure, refer to Supplementary Table 4. Scale bars: 5 µm (B), 10 µm (D).

**Supplementary Figure 11.**
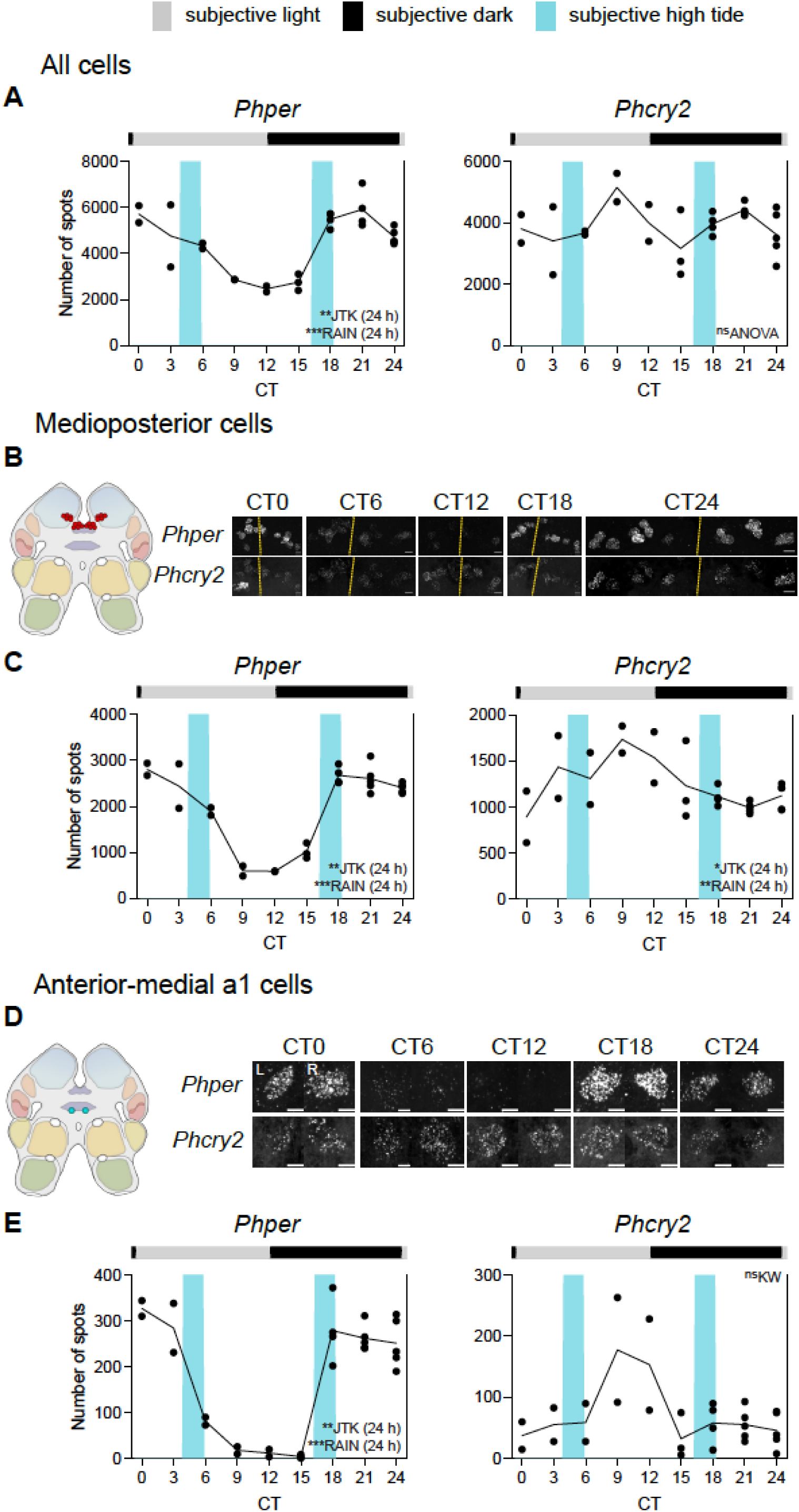
Free-running clock gene expression rhythms in tidally entrained *P. hawaiensis* (tidal phase θ +5.4 h). (A) Mean aggregate expression, quantified as the number of FISH spots, of *Phper* and *Phcry2* across all cells plotted across time in tidally entrained animals sampled under DD. Each point is from one brain. (B) Left: Cartoon to show location of medioposterior cells in the brain and Right: representative maximum intensity Z-projections of *Phper* and *Phcry2* expression in the anterior-medial a1 cells from both hemispheres (L: left, R: right, yellow dashed lines: midlines) across CT (individual HCR-FISH channels in rows). (C) Mean expression, quantified as the number of FISH spots, across CT for *Phper* (left) and *Phcry2* (right) in the anterior-medial a1 cells. Each point is from one brain. (D, E) As in B, C for anterior-medial a1 cell group. Inset text indicates statistical tests performed to determine variation of gene expression with time and/or rhythmicity, with the tested periods (if any) in parentheses. *p* <0.001***, *p* <0.01**, *p* ≤0.05*, *p* >0.05^ns^. KW: Kruskal-Wallis test. For statistics shown in this figure, refer to Supplementary Table 5. Scale bars: 5 µm (B), 10 µm (D).

**Supplementary Figure 12.**
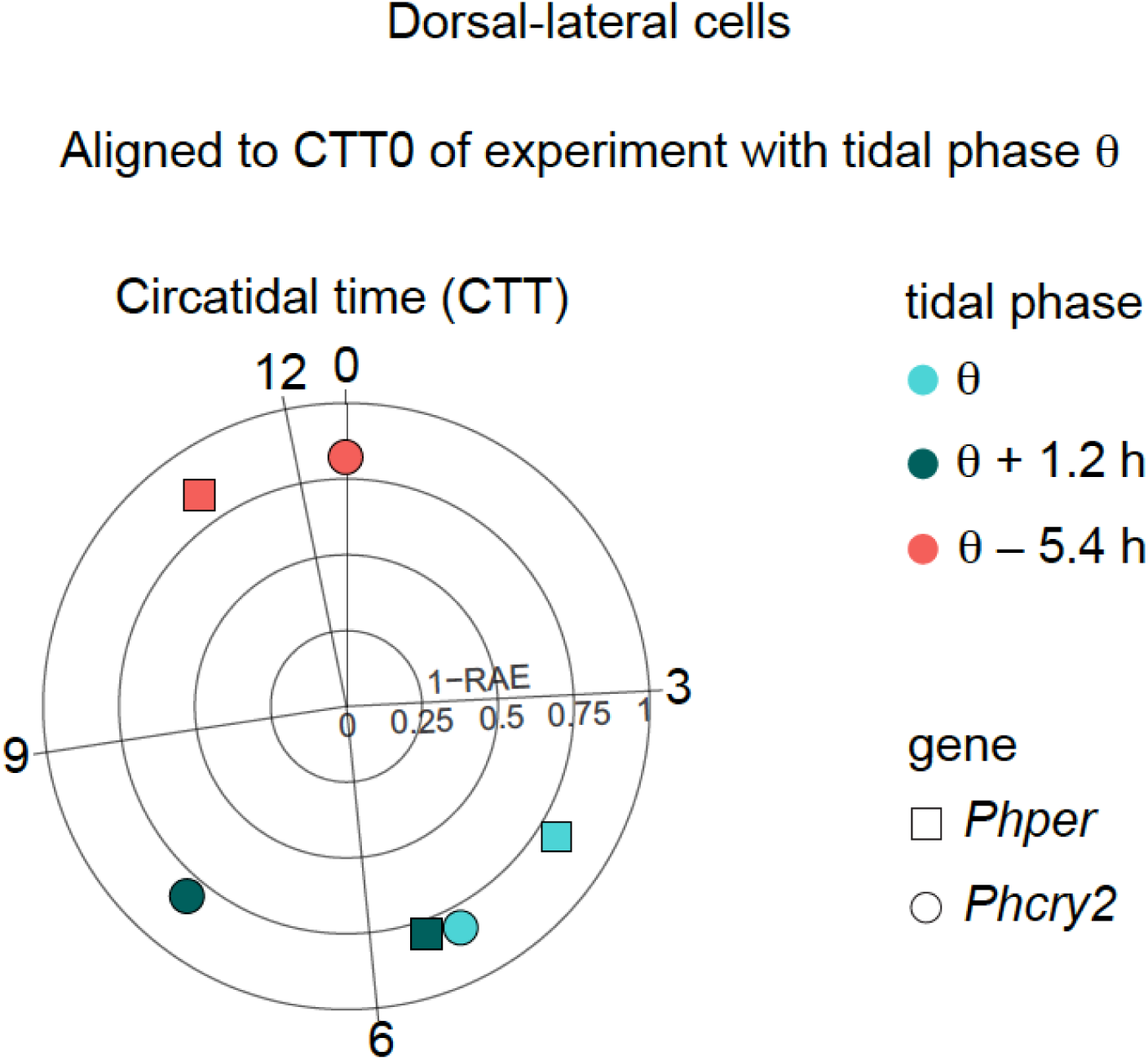
Acrophase of free-running clock gene expression pattern in the dorsal lateral cells of *P. hawaiensis* entrained to tidal cycles of different phase. Circular plots of phase of circatidal peak expression of *Phper* and *Phcry2* in the dorsal-lateral cells of *P. hawaiensis* exhibiting circatidal activity rhythms. The phase of each experiment is aligned to circatidal time (CTT) =0 of the experiment with tidal phase θ. Each colour represents an individual experiment at each of different phases (total N =3 × 3). RAE: relative amplitude error.

**Supplementary Figure 13.**
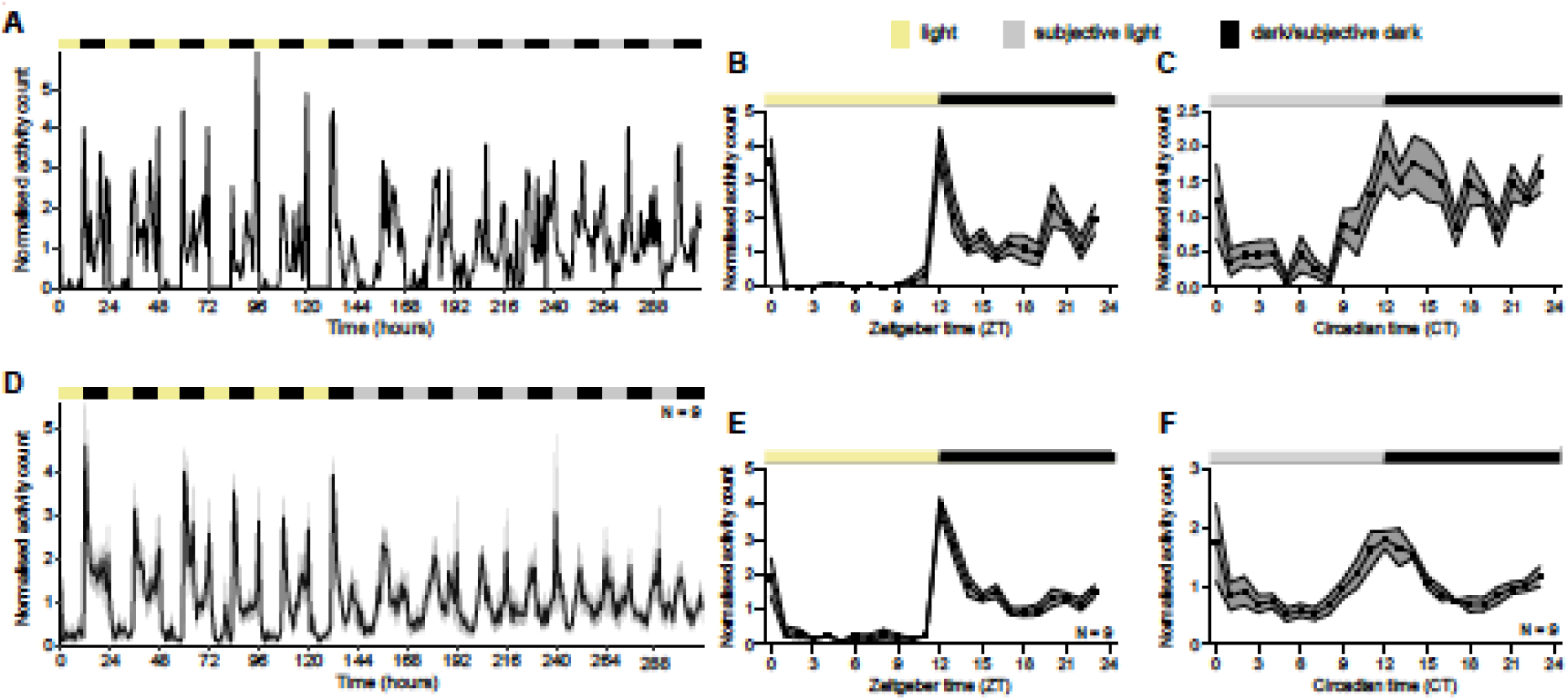
Laboratory-cultured *P. hawaiensis* show circadian activity rhythms after entrainment to LD cycle. (A) Representative activity recording of individual laboratory-cultured *P. hawaiensis*, which had never been subjected to tidal entrainment, initially maintained under 12 h:12 h L:D cycle (yellow/black bars) and then allowed to free-run under DD (grey/ black bars). (B) Mean activity profile for the individual from A under LD showing higher activity during night-time. Shaded area indicates ±SEM. (C) Mean activity profile for the individual from A under DD showing higher activity during subjective night. Shaded area indicates ±SEM. (D, E and F) as for A, B and C but showing group data as mean ±SEM (shading, N =9).

**Supplementary Figure 14.**
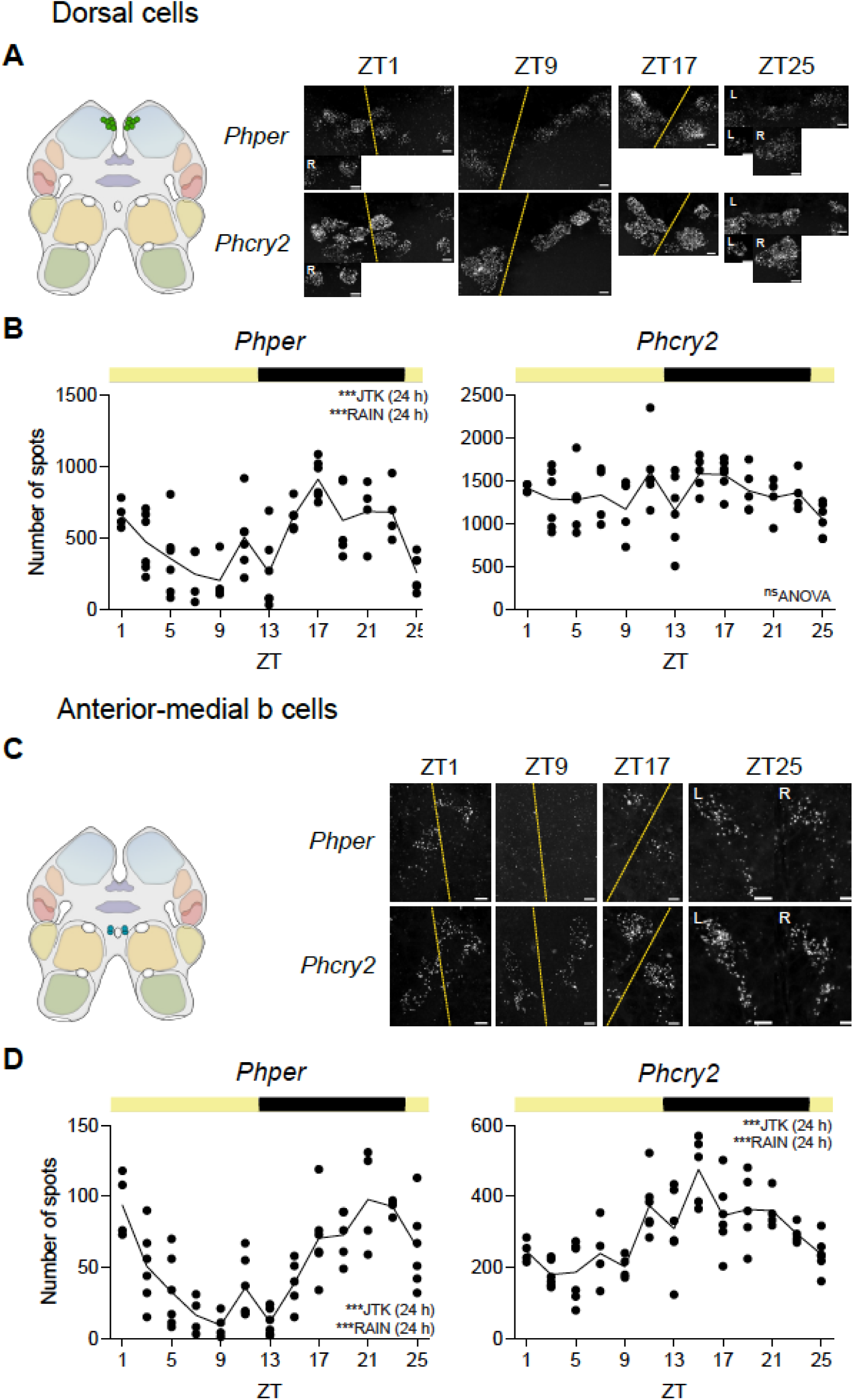
Time course of rhythmic expression of *per* and *cry2* in cells of *P. hawaiensis* synchronised to the LD cycle. (A) Left: Cartoon to show location of medioposterior cell group in the brain and Right: representative maximum intensity Z-projections of *Phper* and *Phcry2* expression in the medioposterior cells of both hemispheres (L: left, R: right, yellow dashed lines: midlines) across ZT (individual HCR-FISH channels in rows). (B) Mean expression, quantified as the number of FISH spots, of *Phper* (left) and *Phcry2* in the dorsal cells plotted across daily time. Each point is from one brain. (C, D) as in A, B for anterior-medial b cell group. Inset text indicate statistical tests performed to determine variation of gene expression with time and/or rhythmicity, with the tested periods (if any) in parentheses. *p* <0.001***, *p* <0.01**, *p* ≤0.05*, *p* >0.05^ns^. Scale bars: 10 µm (b), 5 µm (d, f, h). For statistics shown in this figure, refer to Supplementary Table 6.

**Supplementary Figure 15.**
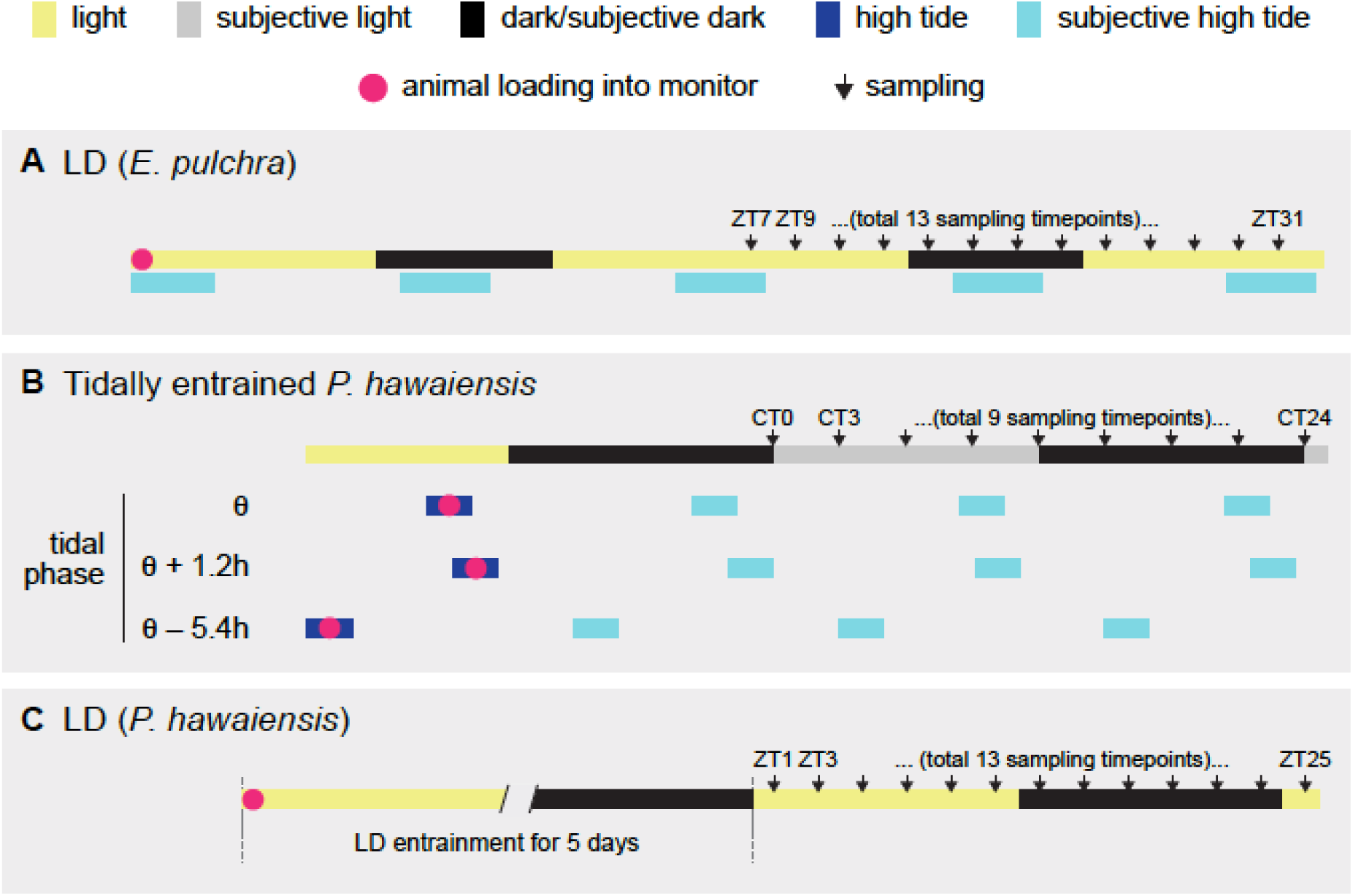
HCR-FISH time-course design. (A) Field-collected *E. pulchra* were loaded into activity monitors and maintained under 16 h: 8 h LD cycles. At ZT7 on the day after loading, animals showing circatidal rhythmicity in their swimming activity were sampled every 2 hours, for a total of 13 timepoints. (B) Laboratory-cultured *P. hawaiensis* which had been subjected to tidal cycles of mechanical agitations were loaded into activity monitors whilst being shaken. At CT0 on the day after loading, animals showing peaks in activity coinciding with the subjective high tide during the night before were sampled every 3 hours, for a total of 9 timepoints. Time-course design is also described in Supplementary Table 10. (C) Laboratory-cultured *P. hawaiensis* that had never been tidally entrained were loaded into activity monitors and maintained under 12 h: 12 h LD cycles for 5 days. Animals showing 24-h rhythmicity over the 5-day LD entrainment period were sampled every 2 hours beginning at ZT1 on day 6, for a total of 13 timepoints. Time-course design is also described in Supplementary Table 10.

**Supplementary Table 1.** Summary of cell groups enriched for *per* and *cry* expression in *E. pulchra* and *P. hawaiensis*.

| ***E. pulchra*** |  |  | ***P. hawaiiensis*** |  |  |
| --- | --- | --- | --- | --- | --- |
| Cell group | # of cells | Anatomical location | Cell group | # of cells | Anatomical location |
| Medioposterior | 18 | rPBp | Medioposterior | 20 | rPBp |
| Medial | 10 | rPBdp | Medial | 6 | rPBp |
| Dorsal | 8 | rHE/MTp | Dorsal | 12 | rHE/MTd |
| Medioanterior | 2 | rMPa | Dorsal-lateral | 4 | rHE/MTd |
| Lateroposterior | 4 | rMTp | Anterior-lateral | 2 | rHE/MTa |
| Dorsoanterior | 4 | rHE/MTa | Anterior-medial a1 | 2 | rMPa |
| Lateroanterior | 2 | rLOva | Anterior-medial a2 | 2 | rMPa |
| Ventroposterior | 8-12 | rMPp | Anterior-medial b | 4 | rMPa |
|  |  |  | Anterior-medial c | 8-10 | rMPa |

Note: because of its location, the ventroposterior cell group in *E. pulchra* could not be imaged in the time-course studies.

**Supplementary Table 2.**
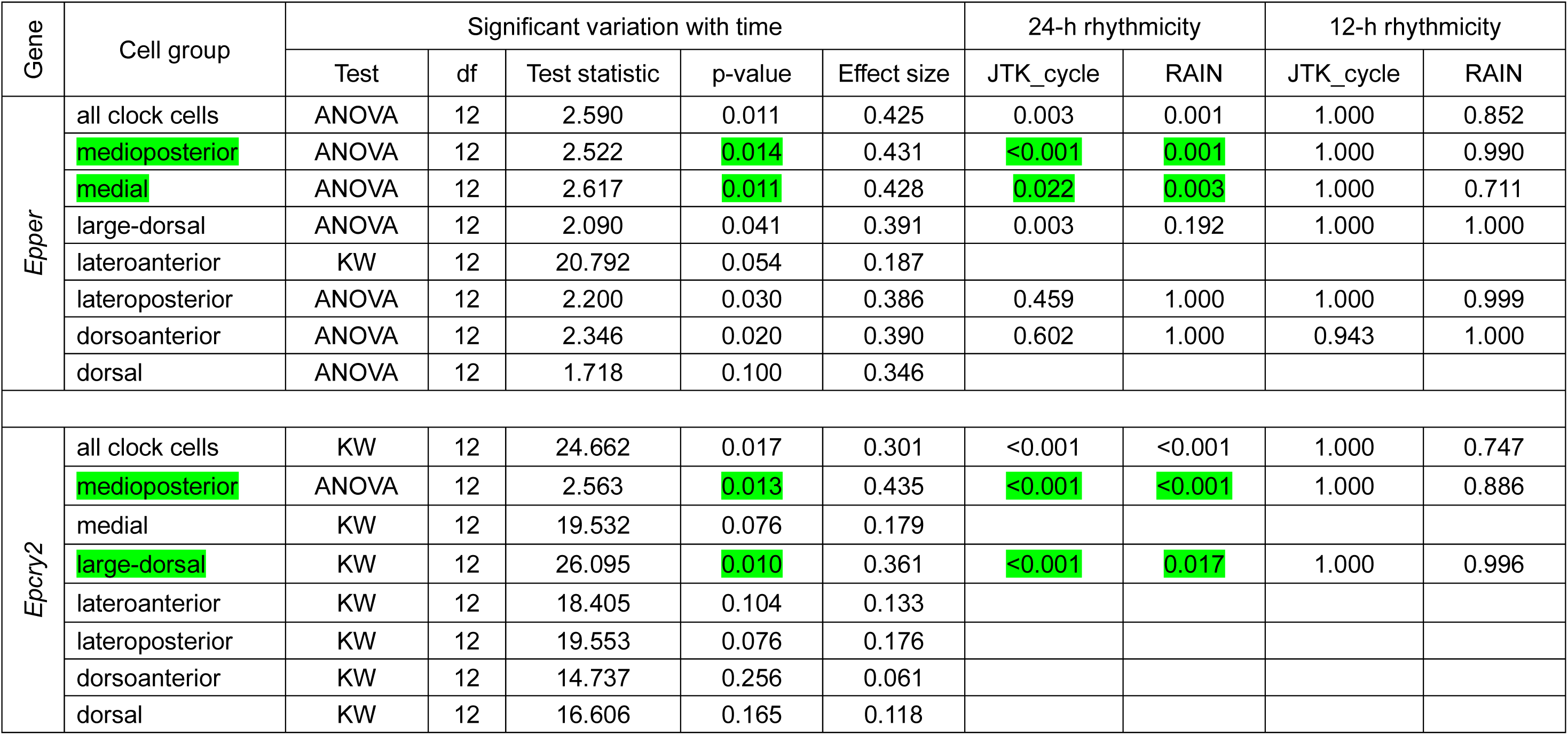
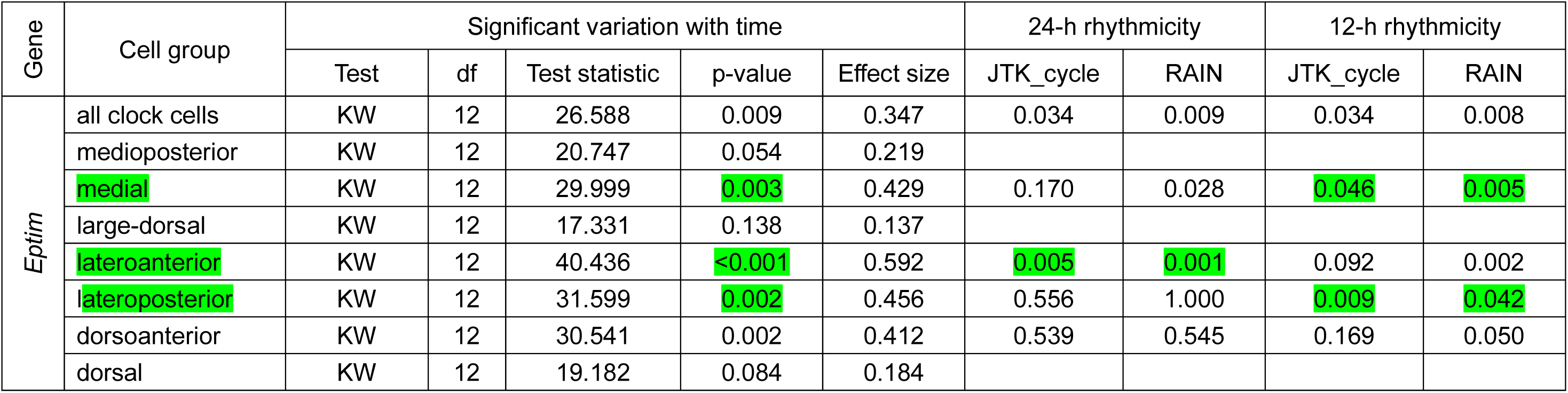
Statistical tests for all gene-cell group combinations for the HCR-FISH time-course experiment of tidally active, beach-collected *E. pulchra* under LD. Effect size quantified as eta squared. Related to Figure 3 and Supplementary Figure 8. KW: Kruskal-Wallis test. Cell groups with statistically significant periodicities.

**Supplementary Table 3.**
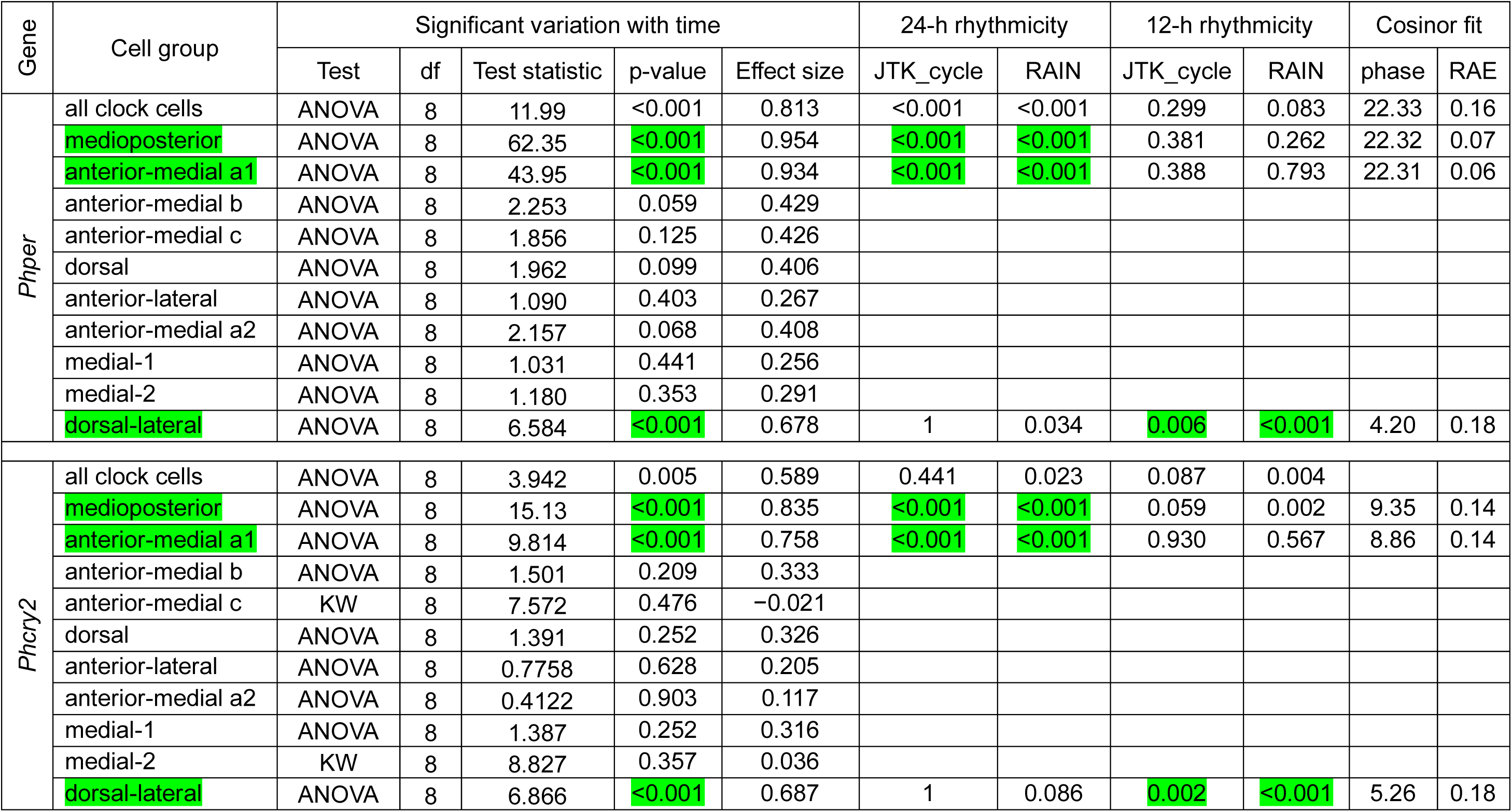
Statistical tests and cosinor analysis (with defined periods of 24-h and/or 12.4-h, whenever appropriate) for all gene-cell group combinations for the HCR-FISH time-course experiment of *P. hawaiensis* entrained to tidal phase θ. Related to Figure 4 and Supplementary Table 7. Effect size quantified as eta squared. Phase derived from cosinor fit is corrected to Circadian Time (CT) =0 or Circatidal Time (CTT) =0, as appropriate. KW: Kruskal-Wallis test; RAE: relative amplitude error. Cell groups with statistically significant periodicities.

**Supplementary Table 4.** Statistical tests and cosinor analysis (with defined periods of 24-h and/ or 12.4-h, whenever appropriate) for all gene-cell group combinations for the HCR-FISH time-course experiment of *Parhyale hawaiensis* entrained to tidal phase θ +1.2 h. Related to Figure 5, Supplementary Figure 10 and Supplementary Table 7. Effect size quantified as eta squared. Phase derived from cosinor fit is corrected to Circadian Time (CT) =0 or Circatidal Time (CTT) =0, as appropriate. KW: Kruskal-Wallis test; RAE: relative amplitude error. Cell groups with statistically significant periodicities.

| Gene | Cell group | Significant variation with time |  |  |  |  | 24-h rhythmicity |  | 12-h rhythmicity |  | Cosinor fit |  |
| --- | --- | --- | --- | --- | --- | --- | --- | --- | --- | --- | --- | --- |
|  |  | Test | df | Test statistic | p-value | Effect size | JTK_cycle | RAIN | JTK_cycle | RAIN | phase | RAE |
| Phper | all clock cells | ANOVA | 8 | 4.899 | 0.003 | 0.697 | <0.001 | <0.001 | 1 | <0.001 | 23.03 | 0.17 |
|  | medioposterior | ANOVA | 8 | 12.96 | <0.001 | 0.845 | <0.001 | <0.001 | 0.772 | 0.462 | 22.90 | 0.11 |
|  | anterior-medial a1 | ANOVA | 8 | 49.29 | <0.001 | 0.952 | <0.001 | <0.001 | 1 | 0.753 | 22.17 | 0.06 |
|  | anterior-medial b | KW | 8 | 16.20 | 0.040 | 0.304 | 0.232 | 0.003 | 0.772 | 0.106 |  |  |
|  | anterior-medial c | ANOVA | 8 | 1.101 | 0.414 | 0.370 |  |  |  |  |  |  |
|  | dorsal | ANOVA | 8 | 0.7518 | 0.648 | 0.273 |  |  |  |  |  |  |
|  | anterior-lateral | ANOVA | 8 | 1.982 | 0.109 | 0.468 |  |  |  |  |  |  |
|  | anterior-medial a2 | ANOVA | 8 | 1.732 | 0.152 | 0.410 |  |  |  |  |  |  |
|  | medial-1 | ANOVA | 8 | 0.5012 | 0.84 | 0.182 |  |  |  |  |  |  |
|  | medial-2 | ANOVA | 8 | 0.4058 | 0.902 | 0.160 |  |  |  |  |  |  |
|  | dorsal-lateral | ANOVA | 8 | 7.212 | <0.001 | 0.743 | 0.232 | 0.003 | 0.016 | <0.001 | 4.53 | 0.21 |
| Phcry2 | all clock cells | ANOVA | 8 | 1.266 | 0.323 | 0.373 |  |  |  |  |  |  |
|  | medioposterior | ANOVA | 8 | 7.967 | <0.001 | 0.770 | 0.005 | <0.001 | 0.023 | 0.001 | 7.01 | 0.24 |
|  | anterior-medial a1 | ANOVA | 8 | 6.807 | <0.001 | 0.731 | <0.001 | <0.001 | 1 | 0.773 | 7.88 | 0.13 |
|  | anterior-medial b | ANOVA | 8 | 2.880 | 0.026 | 0.535 | 0.222 | 0.014 | 1 | 0.460 |  |  |
|  | anterior-medial c | ANOVA | 8 | 0.4578 | 0.867 | 0.196 |  |  |  |  |  |  |
|  | dorsal | ANOVA | 8 | 0.9256 | 0.522 | 0.316 |  |  |  |  |  |  |
|  | anterior-lateral | ANOVA | 8 | 0.5485 | 0.805 | 0.196 |  |  |  |  |  |  |
|  | anterior-medial a2 | ANOVA | 8 | 1.667 | 0.168 | 0.400 |  |  |  |  |  |  |
|  | medial-1 | ANOVA | 8 | 0.6999 | 0.688 | 0.237 |  |  |  |  |  |  |
|  | medial-2 | ANOVA | 8 | 1.522 | 0.222 | 0.417 |  |  |  |  |  |  |
|  | dorsal-lateral | ANOVA | 8 | 4.908 | 0.002 | 0.663 | 1 | 0.215 | 0.016 | <0.001 | 6.58 | 0.18 |

**Supplementary Table 5.** Statistical tests and cosinor analysis (with defined periods of 24-h and/ or 12.4-h, whenever appropriate) for all gene-cell group combinations for the HCR-FISH time-course experiment of *P. hawaiensis* entrained to tidal phase θ – 5.4 h. Related to Figure 5, Supplementary Figure 11 and Supplementary Table 7. Effect size quantified as eta squared. Phase derived from cosinor fit is corrected to Circadian Time (CT) =0 or Circatidal Time (CTT) =0, as appropriate. KW: Kruskal-Wallis test; RAE: relative amplitude error. Cell groups with statistically significant periodicities.

| Ge | Cell group | Significant variation with time |  |  |  |  | 24-h rhythmicity |  | 12-h rhythmicity |  | Cosinor fit |  |
| --- | --- | --- | --- | --- | --- | --- | --- | --- | --- | --- | --- | --- |
|  |  | Test | df | Test statistic | p-value | Effect size | JTK_cycle | RAIN | JTK_cycle | RAIN | phase | RAE |
| Phper | all clock cells | KW | 8 | 19.98 | 0.010 | 0.705 | 0.002 | <0.001 | 0.627 | 0.25 | 22.32 | 0.16 |
|  | medioposterior | KW | 8 | 20.39 | 0.009 | 0.689 | 0.002 | <0.001 | 1 | 0.879 | 22.57 | 0.11 |
|  | anterior-medial a1 | ANOVA | 8 | 23.67 | <0.001 | 0.913 | 0.005 | <0.001 | 0.829 | 0.654 | 22.73 | 0.12 |
|  | anterior-medial b | ANOVA | 8 | 1.912 | 0.121 | 0.096 |  |  |  |  |  |  |
|  | anterior-medial c | KW | 8 | 9.893 | 0.273 | 0.111 |  |  |  |  |  |  |
|  | dorsal | ANOVA | 8 | 1.808 | 0.145 | 0.460 |  |  |  |  |  |  |
|  | anterior-lateral | KW | 8 | 11.99 | 0.152 | 0.234 |  |  |  |  |  |  |
|  | anterior-medial a2 | KW | 8 | 6.759 | 0.563 | -0.073 |  |  |  |  |  |  |
|  | medial-1 | KW | 8 | 11.56 | 0.172 | 0.223 |  |  |  |  |  |  |
|  | medial-2 | KW | 8 | 7.392 | 0.495 | -0.036 |  |  |  |  |  |  |
|  | dorsal-lateral | ANOVA | 8 | 7.469 | <0.001 | 0.768 | 1 | 0.277 | <0.001 | <0.001 | 4.40 | 0.15 |
| Phcry2 | all clock cells | ANOVA | 8 | 1.544 | 0.215 | 0.421 |  |  |  |  |  |  |
|  | medioposterior | ANOVA | 8 | 2.537 | 0.048 | 0.530 | 0.016 | 0.001 | 1 | 0.84 | 9.06 | 0.23 |
|  | anterior-medial a1 | KW | 8 | 10.06 | 0.261 | 0.114 |  |  |  |  |  |  |
|  | anterior-medial b | ANOVA | 8 | 1.129 | 0.391 | 0.334 |  |  |  |  |  |  |
|  | anterior-medial c | ANOVA | 8 | 2.131 | 0.090 | 0.501 |  |  |  |  |  |  |
|  | medial-1 | ANOVA | 8 | 1.497 | 0.234 | 0.428 |  |  |  |  |  |  |
|  | medial-2 | ANOVA | 8 | 0.8941 | 0.542 | 0.296 |  |  |  |  |  |  |
|  | dorsal | KW | 8 | 13.51 | 0.096 | 0.324 |  |  |  |  |  |  |
|  | anterior-lateral | ANOVA | 8 | 4.823 | 0.003 | 0.694 | 1 | 0.203 | <0.001 | <0.001 | 3.93 | 0.17 |
|  | anterior-medial a2 | KW | 8 | 12.26 | 0.14 | 0.251 |  |  |  |  |  |  |
|  | dorsal-lateral | ANOVA | 8 | 8.943 | <0.001 | 0.799 | 0.758 | 0.085 | <0.001 | <0.001 | 5.59 | 0.18 |

**Supplementary Table 6.**
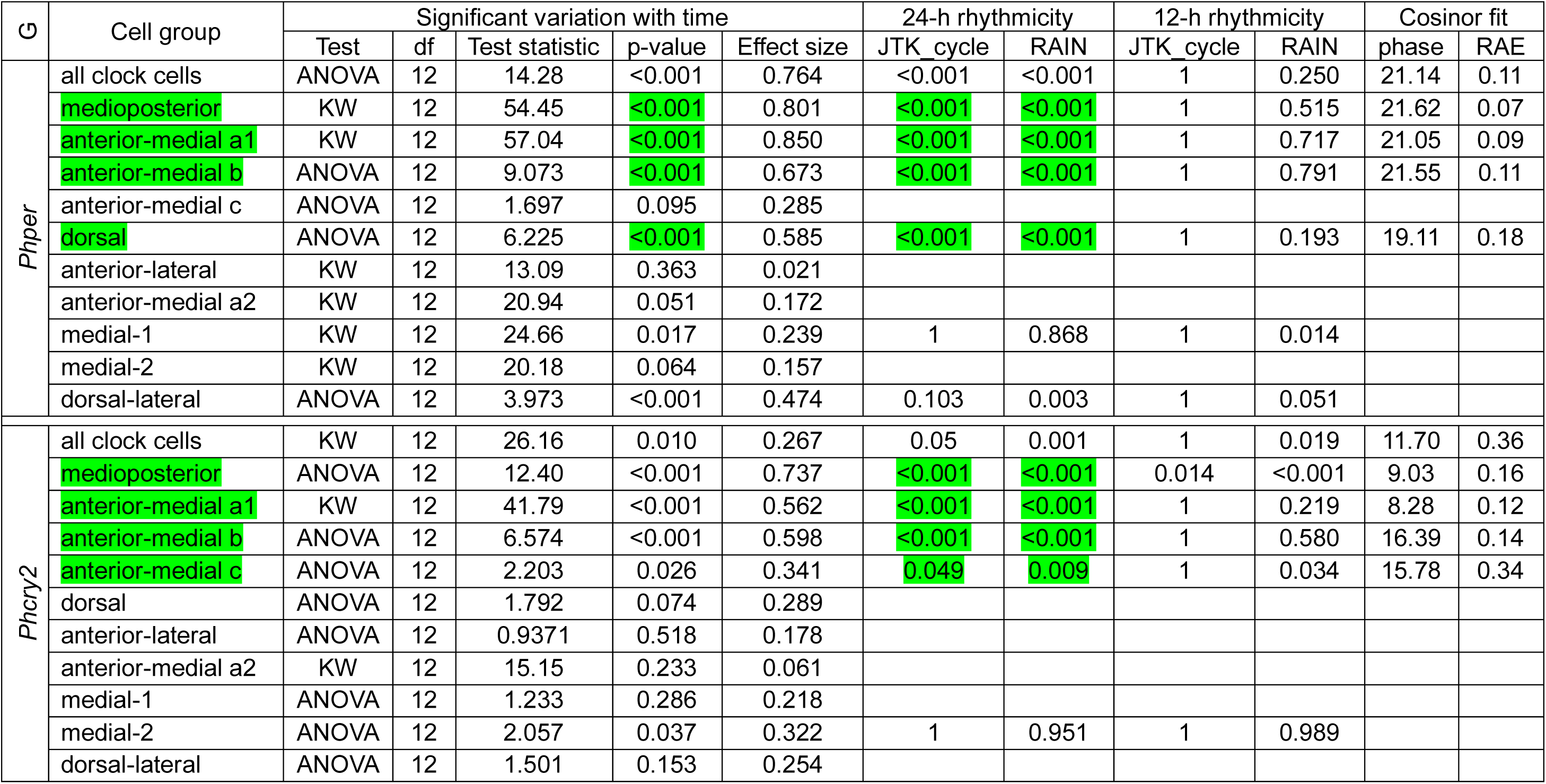
Statistical tests and cosinor analysis (with defined periods of 24-h whenever appropriate) for all gene-cell group combinations for the HCR-FISH time-course experiment of *P. hawaiensis* synchronised to and sampled under LD. Related to Figure 6 and Supplementary Figure 14. Effect size quantified as eta squared. Phase derived from cosinor fit is corrected to Circadian Time (CT) =0. KW: Kruskal-Wallis test; RAE: relative amplitude error. Cell groups with statistically significant periodicities.

| G | Cell group | Significant variation with time |  |  |  |  | 24-h rhythmicity |  | 12-h rhythmicity |  | Cosinor fit |  |
| --- | --- | --- | --- | --- | --- | --- | --- | --- | --- | --- | --- | --- |
|  |  | Test | df | Test statistic | p-value | Effect size | JTK cycle | RAIN | JTK cycle | RAIN | phase | RAE |
| Phper | all clock cells | ANOVA | 12 | 14.28 | <0.001 | 0.764 | <0.001 | <0.001 | 1 | 0.250 | 21.14 | 0.11 |
|  | medioposterior | KW | 12 | 54.45 | <0.001 | 0.801 | <0.001 | <0.001 | 1 | 0.515 | 21.62 | 0.07 |
|  | anterior-medial a1 | KW | 12 | 57.04 | <0.001 | 0.850 | <0.001 | <0.001 | 1 | 0.717 | 21.05 | 0.09 |
|  | anterior-medial b | ANOVA | 12 | 9.073 | <0.001 | 0.673 | <0.001 | <0.001 | 1 | 0.791 | 21.55 | 0.11 |
|  | anterior-medial c | ANOVA | 12 | 1.697 | 0.095 | 0.285 |  |  |  |  |  |  |
|  | dorsal | ANOVA | 12 | 6.225 | <0.001 | 0.585 | <0.001 | <0.001 | 1 | 0.193 | 19.11 | 0.18 |
|  | anterior-lateral | KW | 12 | 13.09 | 0.363 | 0.021 |  |  |  |  |  |  |
|  | anterior-medial a2 | KW | 12 | 20.94 | 0.051 | 0.172 |  |  |  |  |  |  |
|  | medial-1 | KW | 12 | 24.66 | 0.017 | 0.239 | 1 | 0.868 | 1 | 0.014 |  |  |
|  | medial-2 | KW | 12 | 20.18 | 0.064 | 0.157 |  |  |  |  |  |  |
|  | dorsal-lateral | ANOVA | 12 | 3.973 | <0.001 | 0.474 | 0.103 | 0.003 | 1 | 0.051 |  |  |
| Phcry2 | all clock cells | KW | 12 | 26.16 | 0.010 | 0.267 | 0.05 | 0.001 | 1 | 0.019 | 11.70 | 0.36 |
|  | medioposterior | ANOVA | 12 | 12.40 | <0.001 | 0.737 | <0.001 | <0.001 | 0.014 | <0.001 | 9.03 | 0.16 |
|  | anterior-medial a1 | KW | 12 | 41.79 | <0.001 | 0.562 | <0.001 | <0.001 | 1 | 0.219 | 8.28 | 0.12 |
|  | anterior-medial b | ANOVA | 12 | 6.574 | <0.001 | 0.598 | <0.001 | <0.001 | 1 | 0.580 | 16.39 | 0.14 |
|  | anterior-medial c | ANOVA | 12 | 2.203 | 0.026 | 0.341 | 0.049 | 0.009 | 1 | 0.034 | 15.78 | 0.34 |
|  | dorsal | ANOVA | 12 | 1.792 | 0.074 | 0.289 |  |  |  |  |  |  |
|  | anterior-lateral | ANOVA | 12 | 0.9371 | 0.518 | 0.178 |  |  |  |  |  |  |
|  | anterior-medial a2 | KW | 12 | 15.15 | 0.233 | 0.061 |  |  |  |  |  |  |
|  | medial-1 | ANOVA | 12 | 1.233 | 0.286 | 0.218 |  |  |  |  |  |  |
|  | medial-2 | ANOVA | 12 | 2.057 | 0.037 | 0.322 | 1 | 0.951 | 1 | 0.989 |  |  |
|  | dorsal-lateral | ANOVA | 12 | 1.501 | 0.153 | 0.254 |  |  |  |  |  |  |

**Supplementary Table 7.** Circular statistics of gene-cell group combinations that show statistically significant rhythmicity (at 24-h or 12-h) across 3 HCR-FISH time-course experiments of tidally entrained *P. hawaiensis*. Related to Figure 7, Supplementary Figures 10, 11, Supplementary Tables 4-6.

| Gene | Cell group | Mean phase | Mean vector length | Number of experiments |
| --- | --- | --- | --- | --- |
| Circadian oscillations |  |  |  |  |
| *Phper* | All clock cells | 22.56 | 1.00 | 3 |
|  | Medioposterior | 22.60 | 1.00 | 3 |
|  | Anterior-medial a1 | 22.40 | 1.00 | 3 |
| *Phcry2* | Medioposterior | 8.48 | 0.96 | 3 |
|  | Anterior-medial a1 | 8.37 | 0.99 | 2 |
| Circatidal oscillations |  |  |  |  |
| *Phper* | Dorsal-lateral | 4.38 | 1.00 | 3 |
| *Phcry2* |  | 5.81 | 0.96 | 3 |

**Supplementary Table 8.**
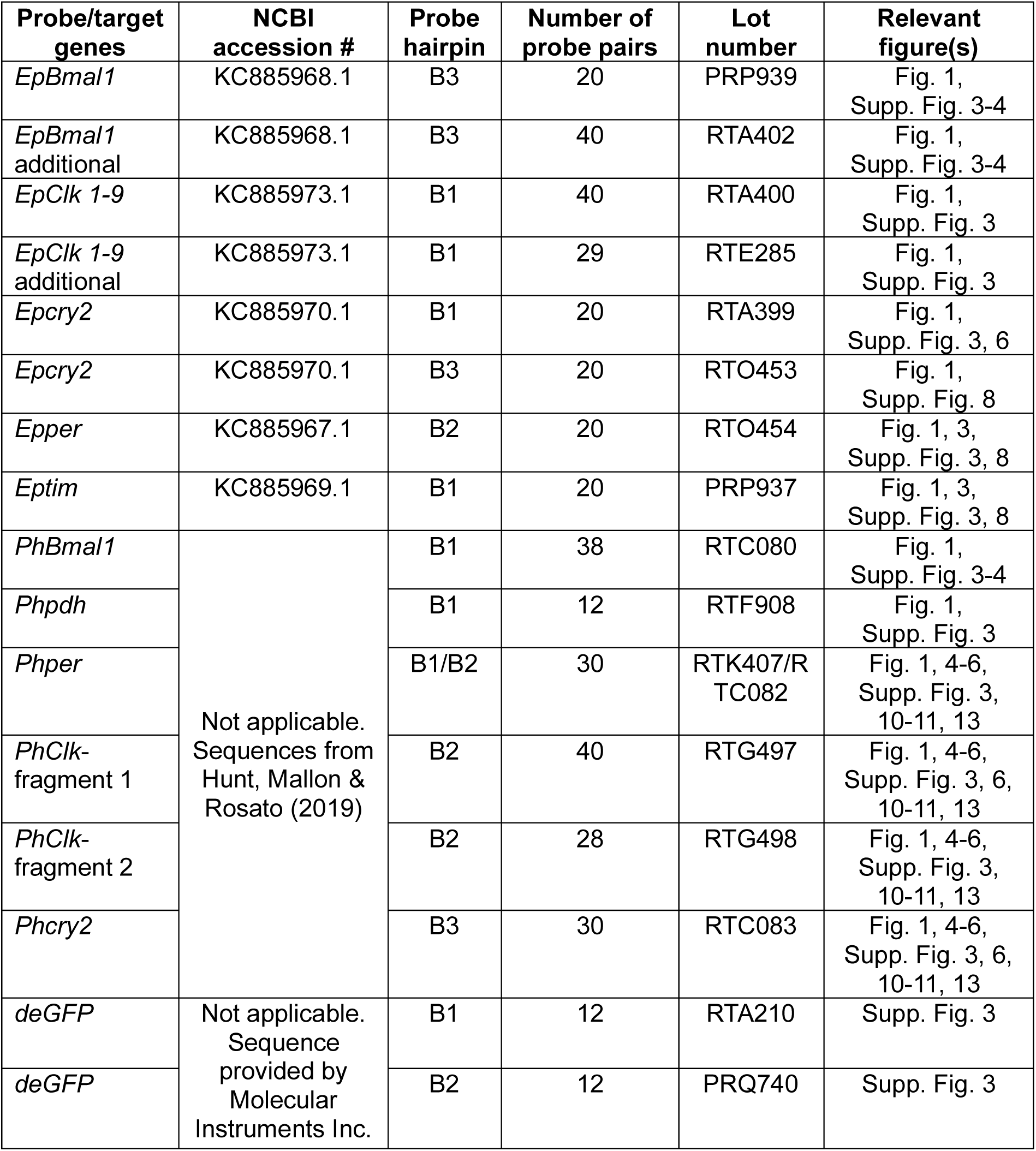
The targets, hairpin identities and lot numbers of the HCR-FISH primary probes used to generate the data described in this study.

| **Probe/target genes** | **NCBI accession #** | **Probe hairpin** | **Number of probe pairs** | **Lot number** | **Relevant figure(s)** |
| --- | --- | --- | --- | --- | --- |
| *EpBmal1* | KC885968.1 | B3 | 20 | PRP939 | Fig. 1, Supp. Fig. 3-4 |
| *EpBmal1* additional | KC885968.1 | B3 | 40 | RTA402 | Fig. 1, Supp. Fig. 3-4 |
| *EpClk 1-9* | KC885973.1 | B1 | 40 | RTA400 | Fig. 1, Supp. Fig. 3 |
| *EpClk 1-9* additional | KC885973.1 | B1 | 29 | RTE285 | Fig. 1, Supp. Fig. 3 |
| *Epcry2* | KC885970.1 | B1 | 20 | RTA399 | Fig. 1, Supp. Fig. 3, 6 |
| *Epcry2* | KC885970.1 | B3 | 20 | RTO453 | Fig. 1, Supp. Fig. 8 |
| *Epper* | KC885967.1 | B2 | 20 | RTO454 | Fig. 1, 3, Supp. Fig. 3, 8 |
| *Eptim* | KC885969.1 | B1 | 20 | PRP937 | Fig. 1, 3, Supp. Fig. 3, 8 |
| *PhBmal1* | Not applicable. Sequences from Hunt, Mallon & Rosato (2019) | B1 | 38 | RTC080 | Fig. 1, Supp. Fig. 3-4 |
| *Phpdh* |  | B1 | 12 | RTF908 | Fig. 1, Supp. Fig. 3 |
| *Phper* |  | B1/B2 | 30 | RTK407/R TC082 | Fig. 1, 4-6, Supp. Fig. 3, 10-11, 13 |
| *PhClk*-fragment 1 |  | B2 | 40 | RTG497 | Fig. 1, 4-6, Supp. Fig. 3, 6, 10-11, 13 |
| *PhClk*-fragment 2 |  | B2 | 28 | RTG498 | Fig. 1, 4-6, Supp. Fig. 3, 10-11, 13 |
| *Phcry2* |  | B3 | 30 | RTC083 | Fig. 1, 4-6, Supp. Fig. 3, 6, 10-11, 13 |
| *deGFP* | Not applicable. Sequence provided by Molecular Instruments Inc. | B1 | 12 | RTA210 | Supp. Fig. 3 |
| *deGFP* |  | B2 | 12 | PRQ740 | Supp. Fig. 3 |

**Supplementary Table 9.** The names, hairpin identities and lot numbers of the RNAScope technology FISH probes.

| Probe/target gene | NCBI accession # | Amplifier | ACD cat number |
| --- | --- | --- | --- |
| *Epper*-C4 | KC885967.1 | C4 | 578211-C4 |
| *EpClk5*-C1 | KC885973.1 | C1 | 578151-C1 |
| *Epcry2*-C1 | KC885970.1 | C1 | 1039411-C1 |
| *Eptim*-C3 | KC885969.1 | C3 | 1099401-C3 |
| *EpBmal1*-C2 | KC885968.1 | C2 | 578191-C2 |
| -ve control |  |  |  |

\* Negative control probe is designed against the DapB gene of *Bacillus subtilis* strain

**Supplementary Table 10.** FISH time-course designs for investigating clock gene expression rhythms. Related to Supplementary Figure 15.

| Experiment | Sampling condition | First sampling time-point | N total animals sampled | Sampling interval | 1<sup>st</sup> subjective high tide onset during sampling | HCR-FISH primary probes | HCR-FISH hairpins (AF: AlexaFluor) | Relevant figures/tables |
| --- | --- | --- | --- | --- | --- | --- | --- | --- |
| 14h:10h LD-maintained *E. pulchra* (Supp. Fig. 15A) |  |  |  |  |  |  |  |  |
| LD (*E. pulchra*) | LD | ZT7 | 65 | 2 | ZT18.4 | *Eptim*-B1, *Epper*-B2, *Epcry2*-B3 | B1-AF647, B2-AF546, B3-AF488 | Fig. 3, Supp. Fig. 8, Supp. Table 2 |
| Tidally entrained *P. hawaiiensis* (Supp. Fig. 15B) |  |  |  |  |  |  |  |  |
| Tidal $\theta$ *P. hawaiiensis* | DD | CT0 | 37 | 3 | CT9.2 | *Phper*-B2, *Phcry2*-B3 | B2-AF546, B3-AF488, | Fig. 4, Supp. Table 3 |
| Tidal $\theta$ + 1.2h *P. hawaiiensis* | DD | CT0 | 35 | 3 | CT10.4 | *Phper*-B2, *Phcry2*-B3 | B2-AF647, B3-AF488, | Fig. 5, Supp. Fig. 10, Supp. Table 4 |
| Tidal $\theta$ – 5.4h *P. hawaiiensis* | DD | CT0 | 35 | 3 | CT3.8 | *Phper*-B2, *Phcry2*-B3 | B2-AF647, B3-AF488, | Fig. 5, Supp. Fig. 11, Supp. Table 5 |
| 12h:12h LD-synchronised *P. hawaiiensis* (Supp. Fig. 15C) |  |  |  |  |  |  |  |  |
| LD (*P. hawaiiensis*) | LD | ZT1 | 79 | 2 | N.A. | *Phper*-B1, *PhClk*-fragment1-B2, *PhClk*-fragment2-B2, *Phcry2*-B3 | B1-AF647, B2-AF546, B4-AF647 | Fig. 6, Supp. Figs. 13, Supp. Table 6 |

## Supplementary Results

### Neuropils of *E. pulchra*

In *E. pulchra*, the optic neuropils comprise lamina (LA), medulla (ME), and lobula (LO). The ME is proximal to the LA and distal to the LO. Ventral to the LO is a small unidentified neuropil. The optic neuropils are distal to the lateral protocerebrum (LP), which comprises the medulla terminalis (MT) and hemi-ellipsoid body (HE). These neuropils are poorly demarcated, and we did not attempt to determine the borders of each, segmenting them as a contiguous complex (HE/MT). The median protocerebrum (MP) comprises a large central body (CB) and protocerebral bridge (PB). The CB extends across the width of the brain and appears to be layered. Ventral to the protocerebrum is the deutocerebrum comprising the deutocerebral chemosensory lobes (DCL) and lateral antenna 1 neuropil (LAN). The DCL are lateral to the brain and comprise numerous olfactory glomeruli. Medial of the DCL is the LAN and ventral of this is antenna 2 neuropil (ANN) within the tritocerebrum. We also identified lateral accessory lobes (LAL) and nodules (NOD), and the medial antenna 1 neuropil (MAN). For *P. hawaiensis* we confirmed the anatomy of Whittforth et al. ^1^ and identified all major neuropils in our refence brain, which correspond closely with those we describe in *E. pulchra*. See also Supplementary Figures 4-5 and Supplementary Videos 1-2.

### Cell groups of E. pulchra and P. hawaiensis

In *E. pulchra, per-* and *cry2*-expressing cells were primarily found in the posterior cell body rind overlying the medial protocerebrum (MP), hemi-ellipsoid body and medulla terminalis (HE/MT, Figure 4a, b). There were nine somata in the rind posterior to the protocerebral bridge (rPBp) ^2^, which we termed the “medioposterior” cells. Dorsal and anterior to these cells was a cluster of 5 somata in the cell body rind dorsal to the PB (rPBd) that we assigned as “medial” cells. Lateral to these medial and medioposterior cells was a cell pair in the rind posterior of the medulla terminalis (rMTp) which we named “lateroposterior” cells. In the cell body rind posterior to the hemi-ellipsoid body/ medulla terminalis (rHEp/MTp) were 4 “dorsal” cells, one of which was larger than the others and we refer to as “large dorsal” cell. In the dorsal region of the cell body rind posterior of the MP (rMPp) ∼4-6 “ventroposterior” cells. In the anterior aspect of the brain, we found fewer cells; in the cell body rind lateroanterior of the hemi-ellipsoid body/medulla terminalis (rHE/MTla) was a single pair of cells, the “dorsoanterior” cells. A single “lateroanterior” cell was in the rind ventral and anterior of the lobula (rLOva). Finally, a pair of cells on the midline (one per hemisphere), anterior of the medial protocerebrum were named “medioanterior” cells (rMPa). Cell groups and their given nomenclature are summarised in Supplementary Table 1.

In *P. hawaiensis*, *per*- and *cry2*-enriched cells were found throughout the anterior and posterior aspects of cell body rind overlying MP and HE/MT. In the cell body rind posterior and lateral to the PB (rPBp) were ten *per-* and *cry2*-enriched somata that we assigned as “medioposterior” cells (Figure 4c, d). Dorsoanterior to these cells were three cells, two of which were closer to each other than the last one; these we termed “medial-2” and “medial-1”, respectively, and collectively, the medial cells. In the cell body rind dorsal of the HE/MT (rHE/MTd), we identified two somata as the “dorsal-lateral” cells, and antero-medial to these cells were six cells that we called the “dorsal” cells. On the anterior aspect of the brain, we identified one cell in the cell body rind anterior of the HE/MT (rHE/MTa) as “anterior-lateral” cells. Lastly, we found 8 – 9 cells on the anterior aspect of the medial protocerebrum (rPMa). We refer to these cells collectively as the “anterior-medial” cells and they can be divided into 4 groups. “Anterior-medial b” cells described two cells with distinctive location lateral to the myoarterial formation. Dorsal to the anterior-medial b cells were two cells, in which one was positioned medially to the other. These, we termed “anterior-medial a1” and “anterior-medial a2” cells, respectively. The “anterior-medial c” cells were a group of 4 – 5 cells located dorsal to the anterior-medial a1-2 cells.

**Supplementary Video 1.**
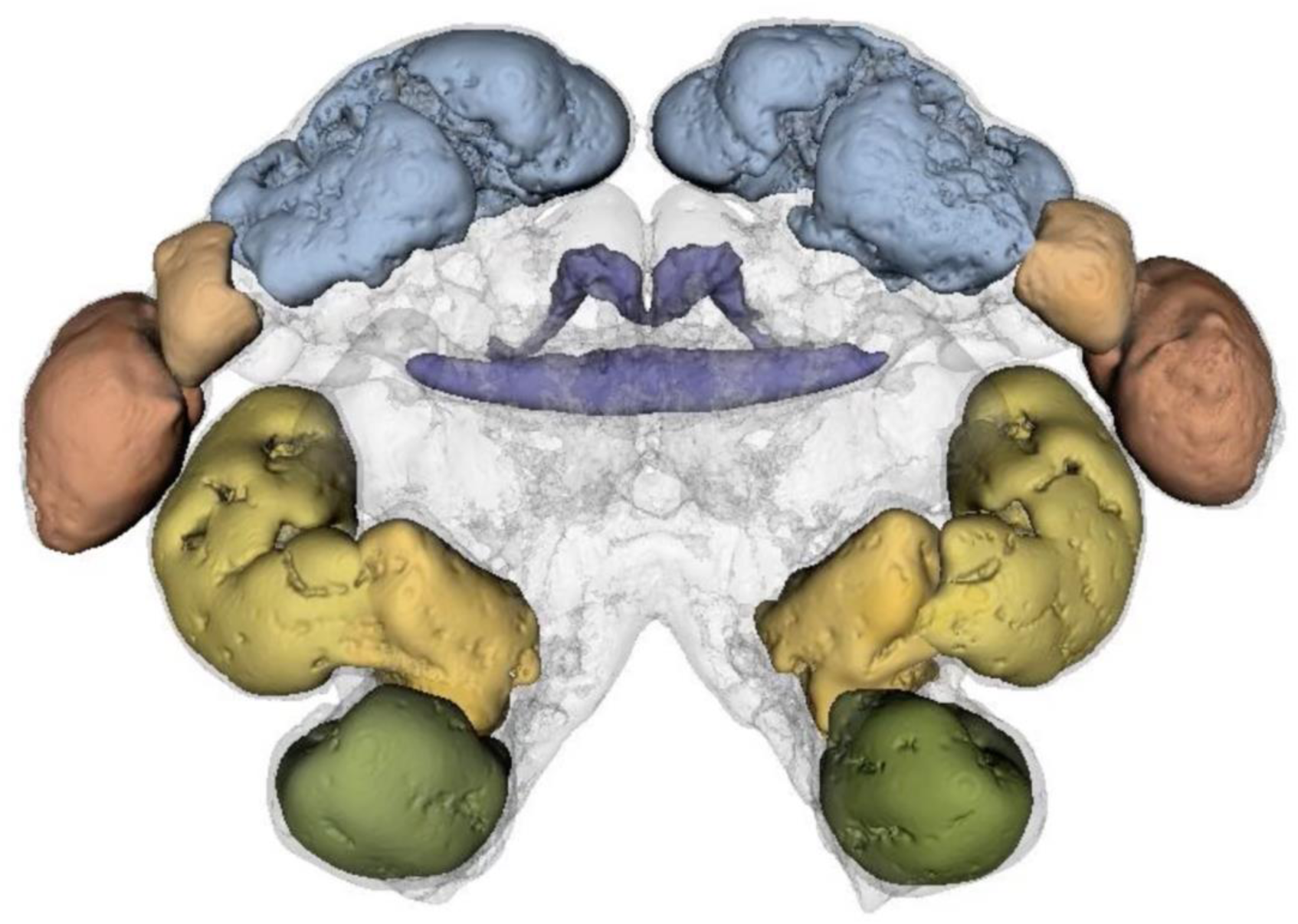
Reference brain of *E. pulchra* revealed by anti-SYNORF1 immunostaining and neuroanatomy of putative clock cell groups revealed by co-immuno-HCR-FISH staining. **Related to Figure 4a and Supplementary Figure 5a.**

**Supplementary Video 2.**
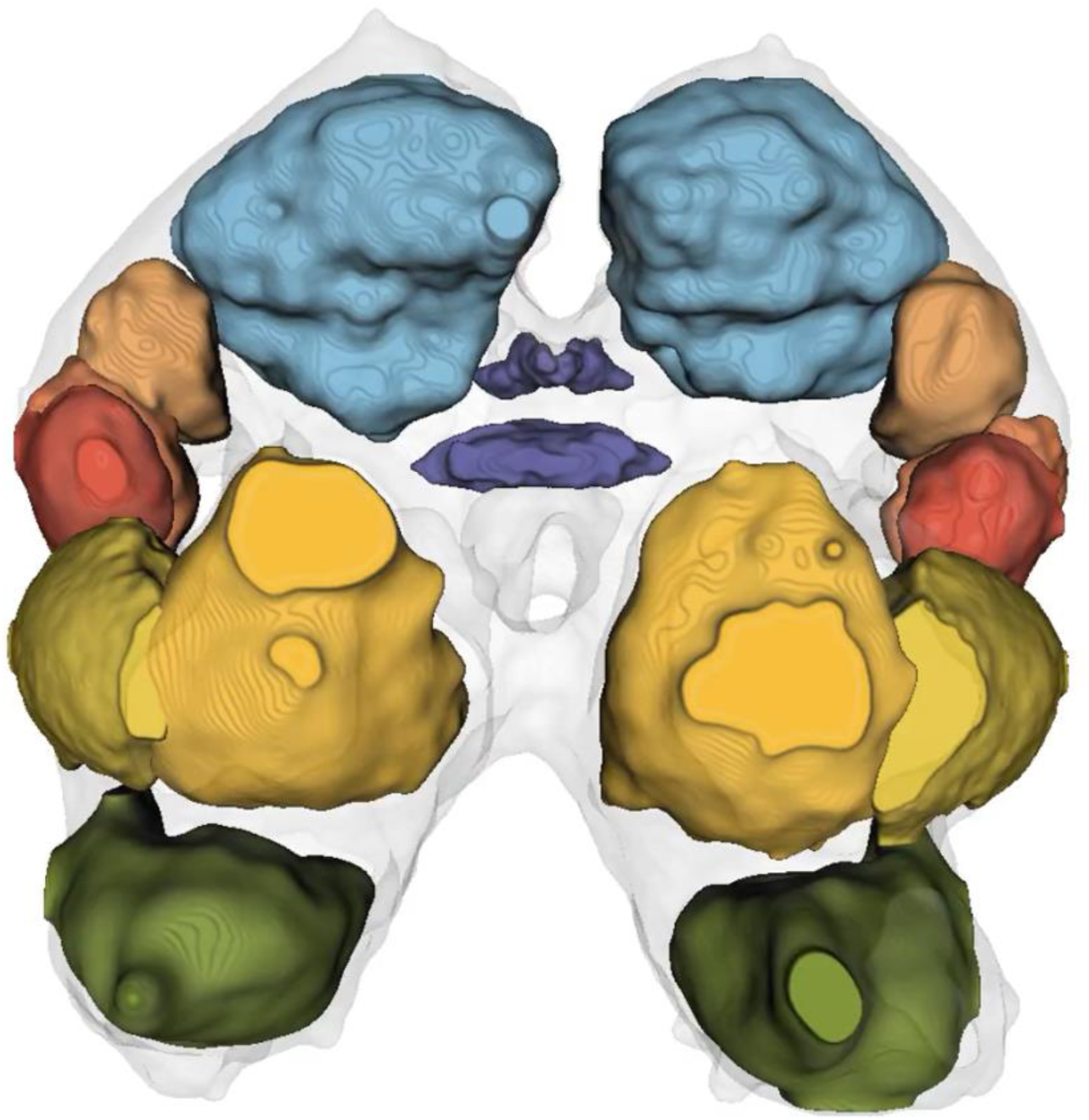
Reference brain of *P. hawaiensis* revealed by anti-SYNORF1 immunostaining and neuroanatomy of putative clock cell groups revealed by co-immuno-HCR-FISH staining. **Related to Figure 4c and Supplementary Figure 5 c.**

## Supplementary methods

### Immunostaining

For immunolabelling of *E. pulchra* brains with anti-SYNORF1, after rehydration to PBST, brains were washed in PTx (PBS supplemented with 0.1% Triton X-100), permeabilised in detergent solution (50 mM Tris-HCl pH 7.5, 150 mM NaCl, 1 mM EDTA pH 8, 1% SDS, 0.5% Tween-20) for 30 min at room temperature (Bruce et al. 2021) and then incubated with 1:50 anti-SYNORF1 in PTx containing 5% normal goat serum (NGS) overnight at room temperature, followed by 5 days at 4°C, with rotation. After extensive washing in PTx, brains were incubated with anti-mouse Alexa 488 (ThermoFisher #A-21141) at 1:1000 in PTx containing 5% NGS for 3 days at 4°C, with rotation. After extensive washing with PTx, nuclei were stained with 0.1 µg/mL DAPI int PTx for 1h. Brains were then bridge-mounted in RapiClear 1.49 (SunJin Lab Optical Clearing Innovation, #RC149001) using No. 2 thickness coverslips to maintain 3D morphology. For *P. hawaiensis*, brains were incubated in blocking buffer (1x PBS, 0.5% Tween-20, 5% NGS, 1% bovine serum albumin) containing 1:50 anti-SYNORF1 for 3 days at 4°C with gentle shaking, followed by extensive washing in PTw (PBS supplemented with 0.5% Tween-20) and incubation in secondary antiserum F(ab’)2-goat anti-mouse IgG-Alexa Fluor 488 (ThermoFisher #A-11017) or Alexa Fluor Plus 555 (ThermoFisher #A48287) at 1:250 made in blocking buffer for 2 days at 4°C with gentle agitation. Nuclei were stained with 0.1 µg/mL DAPI for 30 min prior to final washing in PTw and preparations mounted in DPX (Sigma Aldrich #06522) according to ^3^.

### Reference brains

Whole-mount brain samples used for generating reference brains and describing the neuroanatomy of putative clock cells were imaged on a Nikon CSU-W1 spinning disk confocal microscope equipped with a 25x/1.05NA silicone oil CFI Plan Apochromat Lambda S objective and a Prime 95B 22mm sCMOS camera. To make an *E. pulchra* reference brain, two sex-specific references were made by registering and then averaging 7 male brains to one male “seed” brain and 6 female brains to a female “seed” brain. These sex-specific references were then averaged to generate a final intersex reference brain (an average of 15 brains) used for all subsequent analyses. For *P. hawaiensis*, 10 male brains were registered and then averaged to a seed brain to generate a reference brain. Registration and averaging were done using: https://github.com/jefferislab/MakeAverageBrain ^4 5^. Briefly, Computational Morphometry Toolkit (CMTK) was used to register image stacks against a seed image stack; first with a rigid and then a non-rigid registration ^6,7^. Image stacks of brains with co-immuno-RNA-FISH labelling of neuropils and circadian clock genes (*per*/ *cry2*/ *tim*) were registered to reference brains based on the anti-SYNORF1 signal. Three-dimensional surface models of neuropils were based on reference brains and generated using the Segmentation Editor in 3DSlicer software (www.slicer.org). 3D surface models of putative clock cells were based on circadian clock gene HCR-FISH labelling (*Epcry2* or *Phcry2*) in an image stack registered to the reference brain and using the Segmentation Editor in 3DSlicer.

### HCR-FISH

For HCR-FISH, rehydrated brains were permeabilised in detergent solution for 30 min at room temperature, and equilibrated in pre-warmed probe hybridisation buffer for 30 min at 37°C. Brains were then incubated in probe hybridisation solution containing 10 nM of each primary probe set for 2 days (*E. pulchra*) or 3 days (*P. hawaiensis*) at 37°C. For *P. hawaiensis*, 1:1000 murine RNase inhibitor (NEB #M0314L) was added to the probe hybridisation buffer. Signal amplification with 60 nM fluorescently tagged hairpins was allowed to proceed for 2 days, followed by staining with 0.1 µg/mL DAPI in 5x SSC supplemented with 0.5% Tween-20 (30 min) during one of the final washing steps. *E. pulchra* brains were bridge-mounted using No. 1.5 thickness coverslips in RapiClear 1.49 whilst *P. hawaiensis* brains were dehydrated and mounted in DPX according to ^3^. The target genes used for probe design, their accession numbers, and the Molecular Instruments probe lot numbers and concentrations used are listed in Supplementary Tables 8. For experiments investigating co-expression of clock genes, a 40x/1.3NA Plan Fluorite oil immersion objective was used to image with an optical section interval of 1 µm.

For co-immuno-RNA-FISH staining, following rehydration and permeabilization in detergent solution, *E. pulchra*, brains were incubated with 1:50 anti-SYNORF1 in PTx containing 1.6 U/µL RNaseOUT^TM^ ribonuclease inhibitor (Invitrogen #10777019) overnight at room temperature, followed by 3 days at 4°C, with rotation. After extensive washing in PTx, brains were incubated with anti-mouse Alexa 488 (ThermoFisher #A-21141) at 1:1000 in PTx containing RNase inhibitor for 2 days at 4°C, with rotation. After washing in PTx/w, brains were fixed in 4% PFA (in PBS) for 1 hour at room temperature. After washing in PTx/w, the neuropil-stained brains were processed for HCR-FISH as above and mounted as described above. *P. hawaiensis* brains were washed in PTw and subjected to one-step immunostaining ^8^ by incubating in PTw containing pre-mixed 1:250 murine RNase inhibitor, 1:50 anti-SYNORF1, 3.33 ng/µL Fab goat anti-mouse IgG-Alexa Fluor 488 (Jackson ImmunoResearch #115-547-003) for 2 days at room temperature.

### RNAscope methods for formalin-fixed, paraffin-wax embedded *E.pulchra* brains

*E. pulchra* brains were dissected in ice-chilled crustacean saline ^9^ and fixed immediately in 10% normal buffered formalin (10%) overnight (16h) at RT. After fixing, brains were washed extensively in PBS, dehydrated through an ethanol series (50%, 70%, 80%, 90%, 95%, 100%), cleared in xylene (1h at RT) and infiltrated in molten wax (Leica Paraplast Plus®, Leica Biosystems, Sheffield UK) 3x, each for 1h, before embedding in wax such that the brains could be mounted for frontal sections to be cut. Serial sections (6µm) were cut on a standard wax microtome and mounted on Superfrost™ Plus (Thermo Fisher, UK). After drying, sections were baked at 60°C for 2h to adhere fully to the slides. Slides were de-waxed in clean xylene and taken to water through an ethanol series before drying at 60C for 1h. Following preparation, manufacturer’s instructions were followed exactly, except that target retrieval was done in 700 mL Target Retrieval buffer in a 1L beaker, for 15 minutes at 98-104°C, over a hot-plate. Protease Plus was applied for 30 minutes at 40°C. Signal amplification was done using Opal™ fluors (Opal 520 and Opal 570). All reagents and kit components were as recommended by ACD Biotechne (Abingdon, Oxford, UK. RNAscope® Multiplex Fluorescent Reagent Kit v2 Assay). Target probes were designed by ACD Biotechne and probe details provided in Supplementary Tables 9. Negative control probes were supplied by ACD Biotechne and are designed against the *DapB2* gene of *Bacillus subtilis.* Following hybridisation and amplification, sections were counterstained with DAPI provided with the kit and mounted under Vectashield® Antifade mounting fluid (Vector Laboratories, supplied by 2B Scientific, Oxford, UK). Images were captured on a Leica SP8 confocal microscope at 1µm z-stack intervals and using Leica LAX proprietary software. Images were re-sized, cropped and adjusted for brightness and contrast using ImageJ software.

## Notes

### Competing Interest Statement

The authors have declared no competing interest.

### Summary of Updates

Revised analysis and presentation of the gene expression data with tighter focus on rhythmic cells groups. Consolidation of figures and tables.

